# AAO2 impairment improves aldehyde detoxification by AAO3 in Arabidopsis leaves exposed to UVC or Rose Bengal

**DOI:** 10.1101/2023.09.22.559040

**Authors:** Zhadyrassyn Nurbekova, Sudhakar Srivastava, Nja Zai Du, Veronica Turečková, Miroslav Strand, Rustem Omarov, Moshe Sagi

**Author notes:** The author responsible for distribution of materials integral to the findings presented in this article in accordance with the policy described in the Instructions for Authors (https://academic.oup.com/plphys/pages/General-Instructions) is: Moshe Sagi.

## Abstract

Among the three active aldehyde oxidases in *Arabidopsis thaliana* leaves (AAO1-3), AAO3, which catalyzes the oxidation of abscisic-aldehyde to abscisic-acid, was shown recently to function as a reactive aldehyde detoxifier. Notably, *aao2KO* mutants exhibited less senescence symptoms and lower aldehyde accumulation, such as acrolein, benzaldehyde, and HNE than in wild-type leaves exposed to UV-C or Rose-Bengal. The effect of the absence of AAO2 expression on aldehyde detoxification by AAO3 and/or AAO1 was studied by comparing the response of wild-type plants to the response of *aao1Single* mutant, *aao2KO* mutants and single mutants of *aao3Ss*. Notably, *aao3Ss* exhibited similar aldehyde accumulation and chlorophyll content to *aao2KO* treated with UV-C or Rose-Bengal. In contrast, wild-type and *aao1S* exhibited higher aldehyde accumulation that resulted in lower remaining chlorophyll than in aao*2KO* leaves, indicating that the absence of active AAO2 enhanced AAO3 detoxification activity in *aao2KO* mutants. In support of this notion, employing abscisic-aldehyde as a specific substrate marker for AAO3 activity revealed enhanced AAO3 activity in *aao2KO* and *aao3Ss* leaves compared to wild-type treated with UV-C or Rose Bengal. The similar abscisic acid level accumulated in leaves of unstressed or stressed genotypes indicates that aldehyde detoxification by AAO3 is the cause for better stress resistance in *aao2KO* mutants. Employing the sulfuration process (known to activate aldehyde oxidases) in wild-type, *aao2KO*, and *molybdenum-cofactor sulfurase* (*aba3-1*) mutant plants revealed that the active AAO2 in WT employs sulfuration processes essential for AAO3 activity level, resulting in the lower AAO3 activity in WT than AAO3 activity in *aao2KO*.

## Introduction

Aldehyde oxidase (AO) is a cytoplasmic multicomponent domain enzyme which contains FAD, Fe-S and molybdenum cofactor (MoCo) as its prosthetic groups and thus belongs to the family of Mo-hydroxylases (Koshiba et al. 1996; Yesbergenova et al. 2005). AO catalyzes the oxidation of a variety of aldehydes and heterocyclic compounds, thereby converting them to the respective carboxylic acids (Koshiba et al. 1996, Akaba et al. 1998; Seo et al. 2000a; Seo et al. 2002; Srivastava et al. 2017). AO enzymes have a relatively broad substrate specificity and their role in response to environmental stress factors was previously reported (Omarov et al. 1998; Sagi et al. 1998; Zdunek-Zastocka et al. 2004; Yergaliyev et al. 2016). Early observations demonstrated the presence of four AO genes in *Arabidopsis thaliana* and their organ localization as well as their expression dependency on the developmental stage of the organ. *Arabidopsis* AO1 (AAO1) was shown to be mainly expressed in seeds, roots and seedlings, AAO2 in seedlings and roots, AAO3 in rosettes and roots whereas AAO4 is mainly expressed in siliques (Seo et al. 2000a; Koiwai et al., 2004; Srivastava et al. 2017). Additionally, it was shown that AAO1 is involved in the oxidation of indole-3-carboldehyde (ICHO) to indole-3-carboxylic acid (ICOOH) (Bottcher et al., 2014) and AAO4 is involved in benzoic acid biosynthesis (Ibdah et al. 2009). Additionally, it was demonstrated that AAO4 can efficiently oxidize toxic aldehydes and thus delay silique senescence (Srivastava et al. 2017). Recently it was shown that AAO3 known to oxidize abscisic aldehyde (ABal) to abscisic acid (ABA) (Seo et al. 2000a, 2000b;), can protect rosette leaves from stressed induced aldehyde toxicity (Nurbekova et al. 2021). Additional roles for AAO2 and the other AAOs require further investigation.

Plants are constantly exposed to environmental factors such as extreme temperature, drought, salinity, pathogen attack and natural aging which leads to enhanced generation of aldehydes, downstream to reactive oxygen species (ROS) production. Above a certain level, specific to each of the aldehydes, the elevated level of aldehydes can be toxic, leading to the oxidative injury of plants, enhanced chlorophyll degradation, hastened senescence, and even cell death (Biswas and Mano, 2016; Srivastava et al. 2017; Nurbekova et al. 2021). Detoxification of the increased aldehydes is therefore essential for plant survival and was shown by aldehyde dehydrogenases [ALDH; EC 1.2.1.3 (Sunkar et al. 2003; Stiti et al. 2011; Widhalm and Dudareva, 2015)], aldo-keto reductases (AKR; EC 1.1.1.2), aldehyde reductases [ALR; EC 1.1.1.2 (Oberschall et al. 2000; Yamauchi et al. 2011)], 2-alkenal reductases [AER, EC 1.3.1.74 (Mano et al. 2002)], glutathione transferase tau isozymes [GST (Mano et al. 2019)], and aldehyde oxidases 3 and 4 (Srivastava et al. 2017; Nurbekova et al. 2021).

Recently we observed a significant reduction of AAO2 in-gel activity in plants exposed to UV-C irradiation [Nurbekova et al. 2021; (Fig.7J)], which led us to investigate a possible role of AAO2 in plants exposed to abiotic stresses. In the current study better detoxification of aldehydes in the *aldehyde oxidase 2* knock out (*aao2 KO*) mutants than in WT in response to UV-C irradiation or Rose Bengal application was shown. The application of such stresses resulted in higher chlorophyll degradation in WT leaves, the result of a better oxidation of these aldehydes by *aao2 KO* mutant than in WT. Since *aao2 KO* mutant leaves contains active AAO1 and AAO3, by employing RNA interference (RNAi) on the various T-DNA KO mutants impaired in one of the three aldehyde oxidases AAO1-3, single mutants were generated and were exposed to UV-C irradiation or Rose Bengal application. Notably, independent *aao3* single (*aao3Ss*) mutants containing fully active AAO3 exhibited lower aldehyde accumulation as compared to WT, while *aao1* single (*aao1Ss*) mutants, containing fully active AAO1 resulted in similar or higher level of aldehyde accumulation as well as higher chlorophyll degradation as compared to WT under UV-C irradiation or Rose Bengal application, indicating that the impairment of AAO2 causes the enhancement of AAO3 activity and the capacity of carbonyl aldehyde detoxification in Arabidopsis.

## Results

### Protein expression analysis of *aao2* KO and identification of AAO2 activity

To investigate the possible roles of AAO2 in aldehyde detoxification two independent knock-out (KO) lines of *aao2* mutants [SALK_104895 (KO-95) and SAIL_563_G09 (KO-563)] each carrying exon inserted T-DNA, were employed. The homozygosity of the two *aao2* KO lines were examined. The absence of amplicon when using gene specific left and right primers (LP+RP) and the presence of amplicon with RP and T-DNA specific primers (LB1.3/LB2) indicated the homozygous lines for the knockout mutants of *AAO2* (Fig.1A and Supplemental Table S1). The transcript expression of *AAO2* was determined by quantitative polymerase chain reaction (qPCR) using gene specific primers from either side of T-DNA insertion (Supplemental Tables S2A). No expression was detected in *aao2 KO* mutants, whereas in WT the expression of *AAO2* was evident (Fig.1B) and the amplicons were verified by sequencing (Table *S2B*).

**Figure 1.**
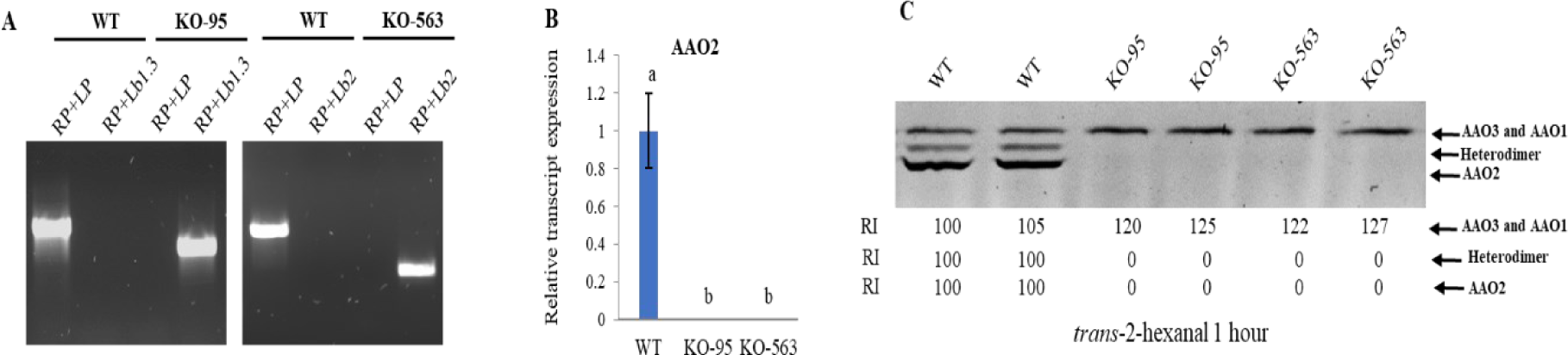
Characterization of *Arabidopsis* aldehyde oxidase 2 mutants. **A.** Verification of AAO2 knock out mutants {*aao2* [SALK_104895 (KO-95) and SAIL_563_G09 (KO-563)]} by PCR using the combination of gene specific primers (LP and RP) and specific primer flanking T-DNA insertion stretches (LB 1.3 or LB2, respectively). **B.** Relative transcript expression of *AAO2* (At3g43600) in WT and the two *aao2* mutants’ rosette leaves. The expression of the *AAO2* in each genotype was compared against *AAO2* in WT after normalization to *Arabidopsis EF-1a* (At5g60390) as the housekeeping gene product and presented as the relative expression. Values means ± SE (n=3). Differences were compared among all the genotypes and different letters show the significant difference between genotypes (Tukey’s honest significant difference, HSD, P value <0.05). **C.** *Arabidopsis* aldehyde oxidases (AAOs) in-gel activity in WT and *aao2* (KO-95 and KO-563). One hundred µg protein extract from three-week old plants’ rosette leaves were fractionated by NATIVE PAGE and AO activity detection with 1 mM *trans*-2-hexanal as the substrate was determined in a reaction solution containing 100 mM Tris-HCl (pH 7.5), 1 mM 3-(4,5-dimethylthiazol-2-yl)-2,5-diphenyltetrazolium bromide and 0.1 mM phenazine methosulfate. The arrow indicates the position of the activity bands generated by AAO1 and AAO3 proteins (uppermost band), middle (heterodimer containing AAO2 with AAO1 and/or with AAO3), and AAO2 (lowermost band). Relative intensity (RI) was estimated by using ImageJ software (https://imagej.nih.gov/ij/) for each activity band. The intensity of the upper, middle, and lowest activity bands in WT was used as reference (100%).

Crude proteins extracted from WT rosette leaves and both *aao2* KOs were employed for the determination of AAO2 in-gel activity by using *trans*-2-hexanal as a substrate (Koshiba et al. 1996; Akaba et al. 1998, 1999; Omarov et al. 1999; Koiwai et al. 2004; Seo et al. 2000a, 2000b; Srivastava et al., 2017). The oxidation of *trans*-2-hexanal led to a highest intensity in the most migrated band and a weaker intensity in the middle band in WT, whereas absence of the middle and the most migrated activity bands was evident in both *aao2* KOs. Notably, the only upper activity band that appeared in *aao2* KOs had a higher band intensity compared to WT (Fig.1C).

To further verify that among the aldehyde oxidases only AAO2 generates the most migrated (lowest) activity band, the lowest activity band in *aao1*, *aao3* KOs and WT (Fig.1C) were excised and peptide sequencing was carried out. The sequencing of trypsinized peptides of the lowest activity band of WT, *aao1* and *aao3* KO mutants revealed 8, 6 and 8 unique peptides of AAO2 respectively (Supplemental Table S3). Additionally, AVSGNLCR and IPTVDTIPK were identified in AAO2 activity band of WT, which share identical sequences in AAO1 and AAO3 respectively. VGGGFGGK peptide of AAO2 that was identified in AAO2 activity band in *aao1KO*, shares identical sequences in AAO1 and AAO3. Similarly, the ASGEPPLLLAASVHCATR peptide of AAO2 identified in *aao3,* shares an identical sequence with AAO3 and therefore in all possible likelihood, it belongs to the AAO2 protein activity band.

### Mutation in AAO2 confers higher capacity of oxidizing aldehydes in *aao2 KO* mutants as compared to WT

The capacity of *aao2* mutant to oxidize a range of aldehydes including reactive aldehydes was compared to WT to get more insight into the function of AAO2. The crude protein extracted from the rosette leaves of *aao2 KO* (KO-95) and WT was fractionated by NATIVE PAGE.

Significantly, knockout of *aao2 KO(KO-95)* mutant exhibited only the upper activity band, whereas WT exhibited additional middle and the lowest activity bands (Fig.1C and Fig.2). The application of various aldehydes at the level of 1 mM [except for HNE (0.25 mM) and ABal (0.1mM)] exhibited that upper in-gel activity band generated by *aao2 (KO-95)* extracted proteins had a more intense activity band (higher activity level) with almost all aldehydes employed as substrates as compared with WT. These results indicate that the absence of AAO2 enhances the oxidation capacity of aldehydes (Fig. 2), suggesting a role for AAO2 in the homeostasis of aldehydes level by modulating another/other aldehyde oxidase/s activity in plant leaves.

**Figure 2.**
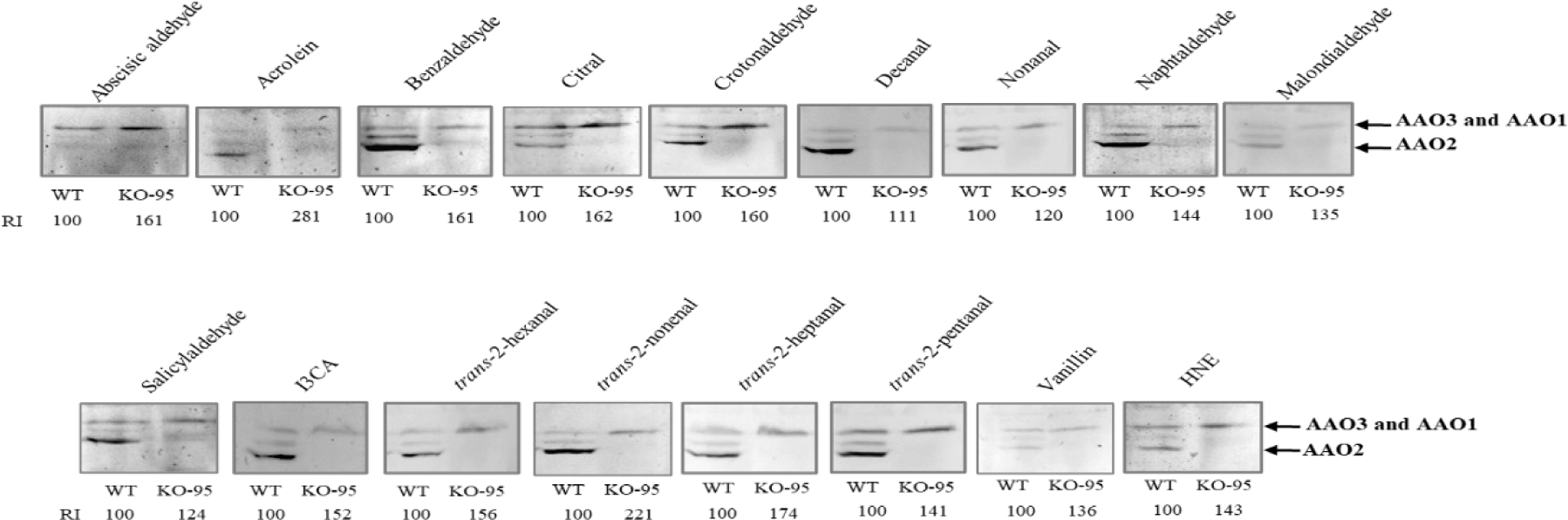
*Arabidopsis* aldehyde oxidases (AAOs) activity assessment in WT and *aao2* [SALK104895 (KO-95)] mutant using different aldehydes as the substrate. One hundred µg protein extracted from rosette leaves of 23-day old WT and KO-95 mutant plants were fractionated by NATIVE PAGE and AO activity with indicated aldehyde as the substrate was carried out. AO enzyme activity was determined in a reaction solution containing 100 mM Tris-HCl (pH 7.5), 1 mM 3-(4,5-dimethylthiazol-2-yl)-2,5-diphenyltetrazolium bromide, 0.1 mM phenazine methosulfate and 1 mM aldehydes, except for HNE and abscisic aldehyde that were applied with 0.25 mM and 0.1 mM, respectively. The reaction was stopped after 3h and after 2 h for abscisic aldehyde, by immersing the gel in 5% acetic acid solution. Thereafter gel images were captured and analyzed for relative intensity (RI) using ImageJ software (http://imagej.nih.gov/ij/) for the slowest (uppermost) migrated band intensity compared to the WT uppermost migrated band as reference (100%).

### UV-C irradiation results-in a faster aldehyde induced senescence in WT compared to *aao2* mutants

The generation of toxic aldehydes which leads to plant senescence is a major response of plants exposed to abiotic stresses. Among the abiotic stresses, UV-C was shown recently to induce premature senescence of siliques and rosette leaves in *Arabidopsis* by enhancing the generation of toxic aldehydes (Srivastava et al. 2017; Nurbekova et al. 2021). Interestingly, a decreased transcript expression of AAO2, as well as declined AAO2 activity but enhanced activity of the middle activity band that contains AAO2 [heterodimer of AAO2 with AAO1 or AAO3 (Akaba et al. 1999; Koiwai et al. 2004)] in leaves was noticed in WT as well as *aao1* and *aao3* mutants exposed to UV-C irradiation (Supplemental Fig. S1). Thus, it was reasonable to study how UV-C irradiation will affect *aao2KO* mutant in the absence of AAO2. Notably, the application of 250 mJ of UV-C irradiation resulted in earlier senescence and greater chlorophyll loss in WT leaves as compared to the two *aao2 KOs (KO-95 and KO-563)* mutants three days after the application of the UV-C irradiation (Fig.3A and B). Determination of the aldehyde level in UV-C treated plants revealed significantly higher levels of acrolein, benzaldehyde and HNE in WT than in the leaves of *aao2 KO* mutant (Fig.3C). An in-gel assay that was carried out to examine the AAO3 and AAO1 activity levels in control (UV-C untreated) and in UV-C treated WT, *aao2KO, aao3KO* (containing active AAO1 and AAO2) and *aao1KO* (containing active AAO3 and AAO2) mutants. Using *trans*-2-nonenal or benzaldehyde as the substrates, revealed enhanced intensity in the most upper (least migrated) activity band in *aao2* mutant compared to WT, *aao1KO* and *aao3KO,* indicating that the absence of active AAO2 enhances the activity of AAO1 and AAO3. The UV-C irradiation resulted in enhancement of the uppermost band in all genotypes tested, indicating the activation of AAO1 and AAO3 in the absence and presence of AAO2 activity. Examination of AAO1 in *aao3* and AAO3 activity in *aao1* mutant revealed that both enzyme activities were enhanced with both aldehyde substrates used, as the result of UV-C irradiation (Fig.3D). Parallel to AAO1 and AAO3 increase, the heterodimer’s activity bands [middle bands (Akaba et al. 1999; Koiwai et al. 2004)] in WT (AAO1:AAO2+AAO3:AAO2) and *aao3* mutant (AAO1:AAO2) exhibited higher increase in response to the applied stress than the heterodimer activity in *aao1* mutant (AAO3:AAO2) with either aldehyde substrate used (Fig.3D). Notably, the AAO3 and/or AAO1 enhanced activity in plants exposed to UV-C irradiation was followed by AAO2 activity (most migrated band) decrease in WT, *aao1KO* and *aao3KO* plants. Accordingly, the absence of AAO2 in *aao2KO* plant resulted in higher activity of AAO3 and/or AAO1, and thus a better aldehyde detoxification and reduced chlorophyll degradation (Fig.3A-D).

**Figure 3.**
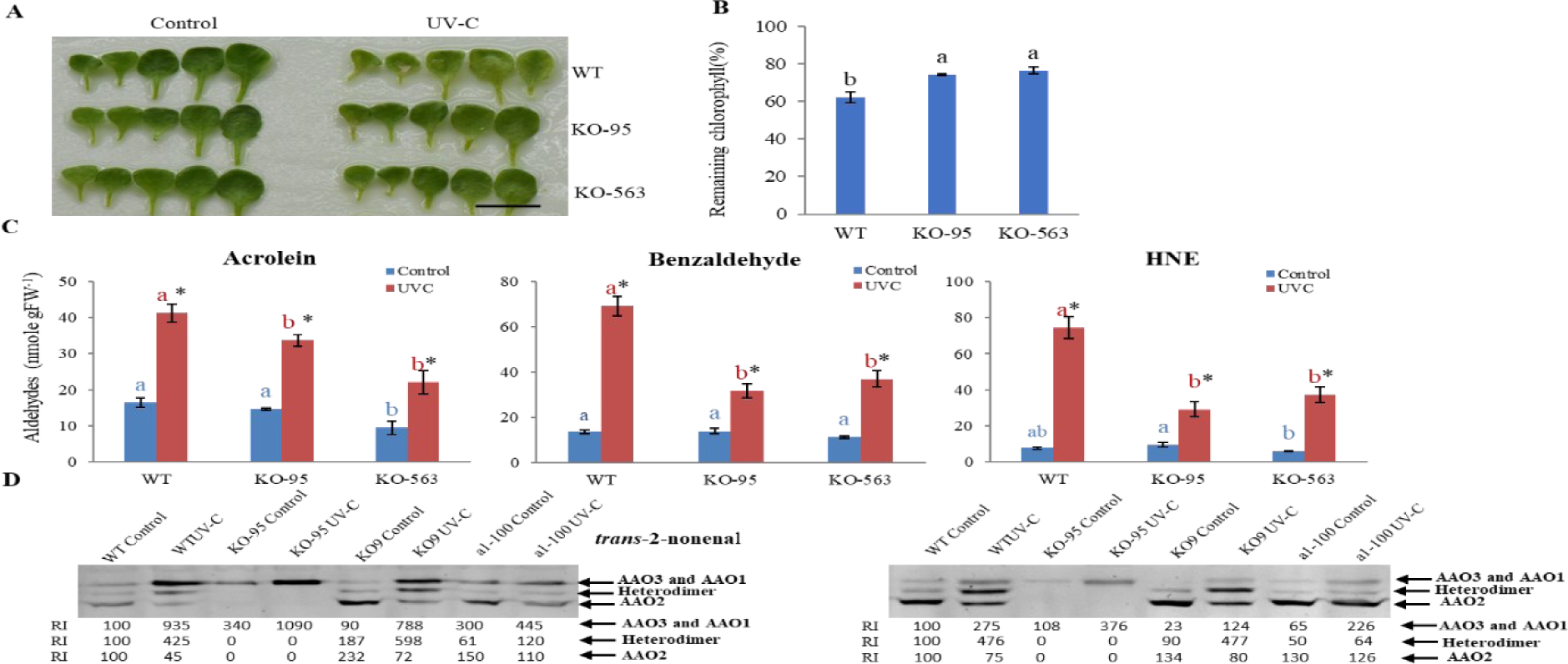
Determination of UV-C-irradiation-induced senescence and senescence-related factors in rosette leaves of *Arabidopsis* wild-type (WT) and *aao2* KO [SALK_104895 (KO-95) and SAIL_563_G09 (KO-563)] mutant plants. **A.** Representative photograph of WT and *aao2* rosette leaves in response to UV-C irradiation. 21-day post germination plants exposed to 250 mJ of UV-C irradiation were kept in a growth room for 72 hours and thereafter documented. Scale bar=1 cm. Rosette leaves were collected 3 days after UV-C treatment and rosette leaves of plants that were not exposed to UV-C were used as the control. First five leaves, oldest to youngest, from left to right are presented. **B.** Remaining chlorophyll in leaves after UV-C treatment. **C.** Indicated aldehyde profiling in control (blue bars) and UV-C treated (red bars) plants. Leaves from 3 different plants were taken as one replica and the bars show the average of at least 4 replicas. Different letters above the bar indicate significant differences (Tukey-Kramer HSD test, P<0.05). Asterisk shows significant differences between treatments within the same genotype (Student’s t test, P < 0.05). **D.** Aldehyde oxidases (AAOs) in gel activity in control and UV-C treated WT, *aao2* (KO-95), *aao3* [SAIL_78_H09 (KO9)] and *aao1* [SALK_018100 (a1-100)]. 150 µg crude protein extracted from rosette leaves were fractionated by NATIVE PAGE and were used for the in-gel activity, assayed for 1h in solution containing 100 mM Tris-HCl (pH 7.5), 1 mM 3-(4,5-dimethylthiazol-2-yl)-2,5-diphenyltetrazolium bromide, 0.1 mM phenazine methosulfate and 1 mM *trans*-2-nonenal or Benzaldehyde as the aldehyde substrate. The gels were scanned, and intensity of the activity bands was estimated using ImageJ software (http://imagej.nih.gov/ij/) and compared with the intensity obtained with UV-C untreated (control) WT (employed as 100%) and presented as relative intensity (RI).

### Application of Rose Bengal confers aldehyde induced earlier senescence in WT compared to *aao2* mutant leaves

Rose Bengal is a well-known singlet oxygen generator (Knox and Dodge, 1984; Havaux and Triantaphylide, 2009) that participate in lipid peroxidation resulting in the generation of toxic aldehydes that above a certain level can lead to programmed cell death (Triantaphylidès et al. 2008). Applying 0.05 mM Rose Bengal by spray, as another abiotic stress type, induced early senescence symptoms in rosette leaves of WT three days after treatment, whereas *aao2* leaves showed much less visible senescence symptoms (Fig.4A). Detection of the remaining chlorophyll revealed a significant chlorophyll degradation in leaves of WT three days after Rose Bengal application, whereas the chlorophyll was less degraded in *aao2* leaves (Fig.4B). Detection of aldehydes 17 h after Rose Bengal application revealed higher enhancement of acrolein, benzaldehyde, propionaldehyde and HNE in WT than in the leaves of the 2 *aao2* mutants (Fig.4C). In-gel AAO activity assay carried out to examine the AAO3 and AAO1 activity levels in the control treated WT and *aao2* mutant using *trans*-2-nonenal or *trans*-2-hexanal as the substrate, revealed higher activity band intensity in the most upper activity band in *aao2* mutant compared to WT. Using *trans*-2-nonenal as the substrate revealed higher AAO3 activity than AAO1 under control conditions, as indicated by the higher intensity in the most upper band in *aao1* compared to *aao3* mutant. In contrast, *trans*-2-hexanal was a preferable substrate for AAO1 activity, as indicated by the higher intensity of the most upper band in *aao3* compared to *aao1* mutant. The application of Rose Bengal resulted in enhancement of the uppermost band in WT and all examined mutants, indicating the activation of AAO1 and/or AAO3 in the absence and presence of AAO2 activity. Examination of AAO1 in *aao3* and AAO3 activity in *aao1* mutant exposed to Rose Bengal revealed that both enzyme activities were enhanced with either aldehyde substrate used (Fig.4D). Interestingly, the heterodimer activity bands in WT (AAO1:AAO2 and AAO3:AAO2) and in *aao3* mutant (AAO1:AAO2) exhibited an increase in response to the applied stress, whereas the heterodimer activity in *aao1* mutant (AAO3:AAO2) tended to show the absence of enhanced intensity with either aldehyde substrate used (Fig.4D). Notably, similar to the UV-C treatment (Fig. 3D), the AAO3 and/or AAO1 enhanced activity in plants exposed to Ros Bengal spray was followed by AAO2 activity decrease in WT, *aao1KO* and *aao3KO* plants. Accordingly, the absence of AAO2 in *aao2KO* plant resulted in higher activity of AAO3 and/or AAO1, and thus a better aldehyde detoxification and reduced chlorophyll degradation (Fig. 4A-D).

**Figure 4.**
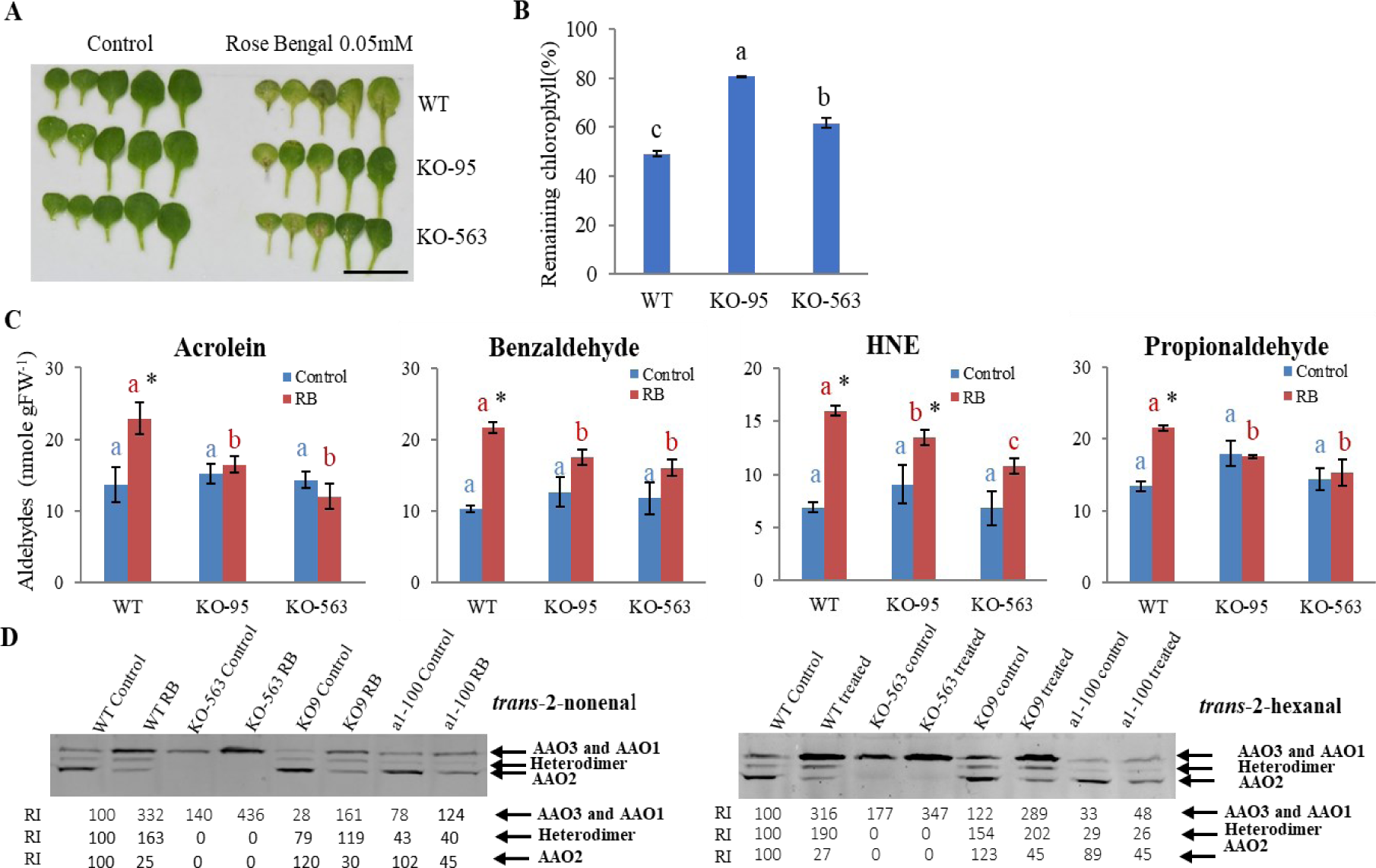
Determination of Rose Bengal-induced senescence and senescence-related factors in rosette leaves of *Arabidopsis* wild-type (WT) and *aao2* KO [SALK_104895 (KO-95) and SAIL_563_G09 (KO-563)] mutant plants. **A.** Representative photograph of WT and *aao2* rosette leaves in response to Rose Bengal application. 23 days post germination plants treated with 0.05 mM Rose Bengal were kept in a growth room for 72 hours and thereafter documented. Scale bar=2 cm. First five leaves, oldest to youngest, from left to right are presented. The rosette leaves were collected 17 hours after Rose Bengal treatment and rosette leaves of plants that were not exposed to Rose Bengal were used as the control. **B.** Remaining chlorophyll in first five leaves after Rose Bengal treatment. **C.** Indicated aldehyde profiling in control (blue bars) and Rose Bengal treated (red bars) plants. Leaves from 3 different plants were taken as one replica and the bars show the average of at least 4 replicas. Different letters above the bar indicate significant differences (Tukey-Kramer HSD test, P<0.05). Asterisk shows significant differences between treatments within the same genotype (Student’s t test, P < 0.05). **D.** Aldehyde oxidases (AAOs) in gel activity in control and Rose Bengal treated WT, *aao2* (KO-95), *aao3* [SAIL_78_H09 (KO9)] and *aao1* [SALK_018100 (a1-100)]. 150 µg crude protein extracted from rosette leaves were fractionated by NATIVE PAGE and were used for the in-gel activity, assayed for 1h in solution containing 100 mM Tris-HCl (pH 7.5), 1 mM 3-(4,5-dimethylthiazol-2-yl)-2,5-diphenyltetrazolium bromide, 0.1 mM phenazine methosulfate and 1 mM *trans*-2-nonenal or *trans*-2-Hexanal as the aldehyde substrate. The gels were scanned, and intensity of the activity bands was estimated using ImageJ software (http://imagej.nih.gov/ij/) and compared with the intensity of the activity band obtained with Rose Bengal untreated (control) WT (employed as 100%) and presented as relative intensity (RI).

### The absence of active AAO2 enhances tolerance to UV-C irradiation and Rose Bengal application by improving the capacity of AAO3 to detoxify aldehydes

The higher resistance of *aao2* KO mutants compared to WT exposed to UV-C irradiation and Rose Bengal application prompted us to examine the possible role of each of the two enzymes: AAO1 and AAO3, in the improved tolerance to the stresses by generating single functioning mutants. The generation of single functioning *aao1* (*aao1S*) was done by silencing *AAO3* in the background of *aao2* (KO-95), while *aao3S* was generated by silencing *AAO1* in the background of *aao2* (KO-95) or *AAO2* silencing in the background of *aao1* KO mutant. The AAO-compromised lines were exposed to UV-C irradiation or Rose Bengal spray after verifications of the mutations by detection of the transcript’s expression of the targeted genes as compared to the expression in WT leaves (Supplemental Fig. S2).

Notably, rosette leaves of the *aao1S (aao1S-11)* mutants impaired in AAO3 and AAO2 expressions exhibited significantly lower remaining chlorophyll level than WT leaves 3 days after exposing to 250 mJ of UV-C irradiation or 0.05 mM of Rose Bengal application. In contrast, *aao3S* mutants *(aao3S-1, aao3S-7, aao3S-12, aao3S-18)* impaired in AAO1 and AAO2 expression, as well as the *aao2* (KO-95) mutant exhibited significantly higher remaining chlorophyll level than WT in response to the applied stresses (Fig.5A, B and Fig.6A, B). Further, detection of aldehydes level in rosette leaves was carried out 3 days after the UV-C irradiation, and revealed significantly higher level of benzaldehyde, crotonaldehyde, propionaldehyde and HNE in WT leaves compared to *aao2 KO* and the three *aao3S (aao3S-1, aao3S-12, aao3S-18)* mutants. In contrast, the levels of these aldehydes in *aao1S* mutant leaves were similar to those in WT (Fig.5C). Detection of aldehydes content in leaves 17h after Rose Bengal application revealed significantly higher accumulation of acrolein, benzaldehyde and HNE in WT and *aao1S* mutant leaves than in leaves of *aao2* (KO-95) and the *aao3S (aao3S-7, aao3S-12, aao3S-18)* mutants (Fig.6C).

**Figure 5.**
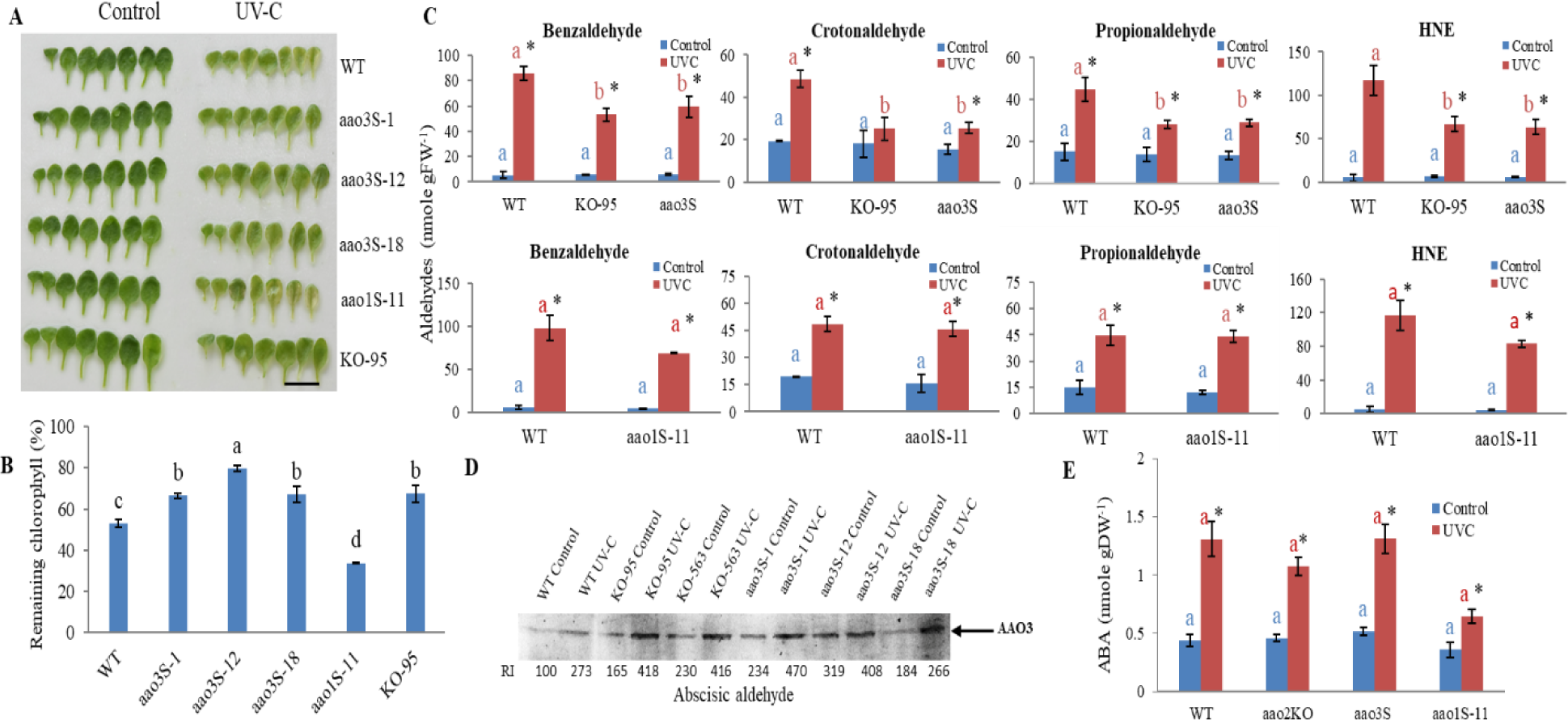
Determination of UV-C-irradiation-induced senescence and senescence-related factors in rosette leaves of *Arabidopsis* aldehyde oxidases single mutants *[aao1Single (aao1S)* and *aao3Singles (aao3S)*], *aao2* KO [SALK_104895 (KO-95) and wild-type (WT) plants. **A.** Representative photograph of WT, *aao2, aao1S* and *aao3S1 (aao3S-1, aao3S-12, aao3S-18)* rosette leaves in untreated (control) and UV-C irradiation treated plants. 21-days post germination (DPG) plants exposed to 250 mJ of UV-C irradiation were kept in a growth room for 72 hours and thereafter documented together with rosette leaves of plants not exposed to UV-C (control). Scale bar=2 cm. **B.** The level of remaining chlorophyll in the first seven leaves (oldest to youngest from left to right) after exposing WT and the various mutant plants to UV-C treatment. **C.** Indicated aldehyde profiling in control (blue bars) and UV-C treated (red bars) plants. Leaves from 3 different plants were taken as one replica and the bars show the average of at least 4 replicas. *aao3S* represents the average of the 3 independent *aao3* single mutants. **D.** Aldehyde oxidase 3 (AAO3) in gel activity in control and UV-C treated WT, *aao2* [KO-95), SAIL_563_G09 (KO-563)] and *aao3S (aao3S-1, aao3S-12, aao3S-18)* using abscisic aldehyde as the specific substrate for AAO3. 150 µg crude protein extracted from WT, *aao2* (KO-95 and KO-563) and *aao3S (aao3S-1, aao3S-12, aao3S-18)* rosette leaves were fractionated by NATIVE PAGE and were used for activity. The gels were scanned after 30 min, and intensity of the activity bands was estimated using ImageJ software (http://imagej.nih.gov/ij/) and compared with that obtained with UV-C untreated (control) WT (employed as 100%) and presented as relative intensity (RI). **E.** Abscisic acid (ABA) level in rosette leaves of WT, *aao2KO (KO-95, KO-563), aao1S (aao1S-11)* and *aao3S* (*aao3S-1, aao3S-12, aao3S-18*) untreated control (blue bars) and UV-C treated (red bars) plants. Different letters above the bar indicate significant difference (Tukey-Kramer HSD test, P<0.05). Asterisk shows significant differences between treatments within the same genotype (Student’s t test, P < 0.05).

**Figure 6.**
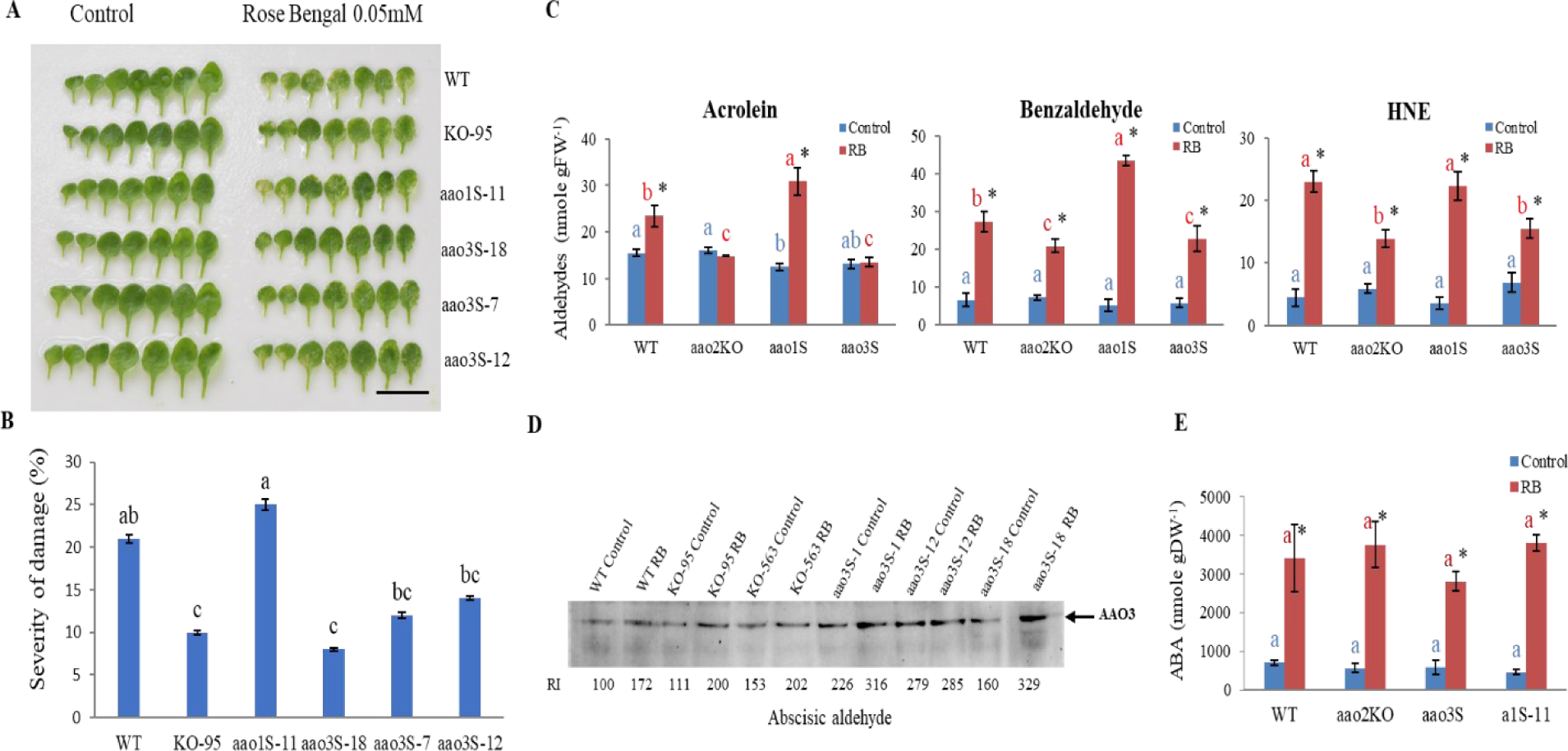
Determination of Rose Bengal-induced senescence and senescence-related factors in rosette leaves of *Arabidopsis* aldehyde oxidases single mutants *[aao1Single (aao1S)* and *aao3Singles (aao3S)*], *aao2* KO [SALK_104895 (KO-95) and wild-type (WT) plants. **A.** Representative photograph of WT, *aao2, aao1S* and *aao3S2 (aao3S-7, aao3S-12, aao3S-18)* rosette leaves in untreated (control) and Rose Bengal treated plants. 21-DPG plants treated with 0.05 mM of Rose Bengal were kept in a growth room for 72 hours and thereafter documented. Scale bar=2 cm. **B.** Damage level in leaves as shown in (A) was calculated as described in the materials and methods. Means +-SEM (n = 6). **C.** Indicated aldehyde profiling in control (blue bars) and Rose Bengal treated (red bars) plants. Seventeen hours after Rose Bengal treatment leaves from 3 different untreated (control) or treated plants were taken as one replica and the bars show the average of at least 4 replicas. *aao3S* is average of the 3 independent *aao3* singles plants. **D.** Aldehyde oxidases 3 (AAO3) in gel activity in control and Rose Bengal treated WT, *aao2* [KO-95), SAIL_563_G09 (KO-563)] and *aao3S* using abscisic aldehyde as the specific substrate for AAO3. 150 µg crude protein extract from WT, *aao2 (*KO-95, KO-563) and *aao3S* rosette leaves was fractionated by NATIVE PAGE and were used for the in-gel activity. The gels were scanned after 1h, and the intensity of the activity bands was estimated using ImageJ software (http://imagej.nih.gov/ij/) and compared with that obtained with Rose Bengal untreated (control) WT (employed as 100%) and presented as relative intensity (RI). **E.** ABA content in rosette leaves of WT, *aao2KO (KO-95, KO-563), aao1S (aao1S-11)* and *aao3S* (*aao3S-7, aao3S-12, aao3S-18*) untreated control (blue bars) and Rose Bengal treated (red bars). Different letters above the bar indicate significant differences (Tukey-Kramer HSD test, P<0.05). Asterisk shows significant differences between treatments within the same genotype (Student’s t test, P < 0.05).

To further rule out the possible role of AAO1 in *aao2* resistance to the applied stresses, *AAO1* overexpression mutant (*A1OE*) used by us before [Nurbekova et al. 2021 (Fig.1B and C)], was exposed together with WT plants to UV-C irradiation that revealed a higher chlorophyll degradation in A1OE (Supplemental Fig. S3A and B). Additionally, similar levels of benzaldehyde, crotonaldehyde, HNE and higher levels of propionaldehyde in *A1OE* compared to WT were detected (Fig. 3C). The results show better resistance of *aao2* KOs and *aao3S* mutants than WT, whereas *aao1S* and *A1OE* mutants exhibited similar resistance to WT when exposed to the abiotic stresses applied (Figs 5A-C, 6A-C, Supplemental Fig. S3). This indicates that AAO1 did not play a role in stress improved resistance of *aao2 KO* as compared to WT while AAO3 is the major player in the detoxification of reactive aldehydes in the absence of active AAO2 in *aao2 KOs*. Additional support to this notion was demonstrated by employing ABal as the indicator substrate specific for AAO3 In-gel activity (Akaba et al. 1998, 1999; Koiwai et al. 2004; Koshiba et al. 1996; Omarov et al. 1999; Seo et al. 2000a,b; Srivastava et al. 2017; Nurbekova et al. 2021) that exhibited higher AAO3 activity level in response to UV-C or Rose Bengal application in all the included genotypes, being significantly higher than WT in *aao2 KO* and *aao3S* mutants. This result indicates that AAO3 is more active in the absence of active AAO2 in *aao2 KO* and *aao3S* mutants compared to WT under the stresses applied (Fig.5D and Fig.6D). Further detection of the ABA level revealed similar enhancement levels in WT compared to *aao2 KO*, *aao3S* and *aao1S* mutants exposed to UV-C irradiation or Rose Bengal application, supporting the notion that the AAO3 - enhanced detoxification activity of aldehyde, rather than ABA level played a role in *aao2 KO* better resistance than in WT and *aao1* mutant (Fig.5E and Fig.6E).

To examine if indeed *aao2KO* mutants can better resist toxic levels of aldehydes than WT, plants were exposed to 4.5 mM benzaldehyde as done recently (Nurbekova et al. 2021) and revealed higher chlorophyll degradation in WT than in the two independent mutants’ leaves 72 h after aldehyde application (Supplemental Fig. S4A and B). These results further indicate the higher capacity of *aao2* KO mutants than WT to protect leaves from aldehyde induced chlorophyll degradation.

## DISCUSSION

### The absence of active AAO2 improves AAO3’s capacity to protect plants from UV-C and RB induced chlorophyll degradation

Early observations showed the presence of three AAO genes in leaves and their expression dependency on the developmental stage and applied abiotic stresses. *Arabidopsis* AAO1 was shown to be mainly expressed in seedlings and rosette leaves (Seo et al. 2000a; Koiwai et al. 2004), oxidizing indole-3-carboldehyde (ICHO) to indole-3-carboxylic acid [ICOOH ((Bottcher et al. 2014))]. AAO3 was shown to be expressed in seedlings and rosette leaves (Seo et al. 2000a; Koiwai et al. 2004), oxidizing abscisic aldehyde (ABal) to abscisic acid (ABA) (Seo et al. 2000a, 2000b;) as well as protecting rosette leaves from stress induced aldehyde toxicity by oxidizing the toxic aldehydes (Nurbekova et al. 2021). The roles of AAO2 are not known and thus require further study. In the current study it was shown that exposing WT and *aao2* mutants, as well as *aao1S* and *aao3S* mutants to UV-C irradiation or Rose Bengal spray revealed enhanced senescence induced by endogenously generated higher level of benzaldehyde, HNE and acrolein in WT and *aao1S* compared to *aao3S and aao2 KOs* mutant’s leaves (Fig.3A-C, Fig.4A-C, 5A-C and 6A-C). These results indicate the essentiality of the absence of active AAO2 and the presence of active AAO3, whereas the presence of active AAO1 is not essential for the better stress resistance genotypes, the *aao2KO*, and the single mutants of *aao3S* under applied stress conditions. The lower remaining chlorophyll content and higher level of propinaldehyde and similar enhacement in benzaldehyde, crotonaldehyde and HNE levels in AAO1OE compared with WT upon UV-C irradiation, further ruled out a possible involvement of AAO1 in *aao2* KO better resistance than WT (Supplemental Fig. S3). Employing In-gel AO assay with ABal as a specific substrate marker for AAO3 activity showed higher AAO3 activity in *aao2 KOs* and *aao3Ss* as compared to WT leaves treated with UV-C or Rose Bengal (Fig.5D and Fig.6D), indicating that enhanced AAO3 capacity plays a role in *aao2KOs* and *aao3Ss* better resistance.

The involvement of reactive aldehydes in stress-induced damage to plants was previously reported, and among lipid peroxide derived carbonyls, α, β-unsaturated aldehydes such as HNE, acrolein and crotonaldehyde were identified as the most reactive species (Yamauchi et al. 2008; Mano et al. 2010, Mano et al. 2014; Srivastava et al. 2017). Interestingly, higher levels of benzaldehyde, acrolein, propionaldehyde and crotonaldehyde were shown to induce early senescence in *aao3* leaves when exposed to UV-C irradiation. Similar to the experimental condition in the current study, *aao3* plants, known as ABA deficient mutants, were grown on half-strength MS media in agar plates sealed with surgical tapes, to control any drought involvement in senescence level in WT and *aao3* mutant (Nurbekova et al., 2021). Indeed, under such controlled condition no effect of ABA or drought stress related was noticed in the *aao3* plants (Nurbekova et al. 2021), whereas similar level of ABA was evident in WT, *aao2KOs, aao1S* and *aao3Ss* in control and stressed plants (Fig.5E and Fig.6E). To support the levels of ABA detected in the stressed plants the expression of *NCED3-2* gene was examined in the leaves of the control and the UV-C or Rose Bengal stressed plants. Considering that *NCED3-2* gene product acts as the commited step of ABA biosynthesis (Ruggiero et al. 2020) generating xanthoxin that leads to the biosynthesis of ABAld, the substrate of AAO3 (Sussmilch et al. 2017), the similar transcript expression of *NCED3-2* evident in WT and *aao2KOs* and *aao3Ss* exposed to the applied stresses (Supplemental Fig. S5), is in agreement with the similar levels of ABA detected in the stressed genotypes. These results indicate that not the ABA level but rather the activated AAO3 is reponsible for the higher level of the remaining chlorophyll by better aldehyde detoxification capacity in *aao2KOs* and *aao3Ss* mutant plants exposed to UV-C irradiation or Rose Bengal application (Fig.5D and Fig.6D).

### The absence of active AAO2 in *aao2 KO* mutants enhances AAO3 activity as the result of improved sulfuration by ABA3

The decrease in AAO2 activity in WT as well as the *aao1* and *aao3* mutants exposed to UV-C or Rose Bengal (Figs. 3D, 4D), and specifically the higher AAO3 activity than in WT, in the mutants absent active AAO2, the *aao2 KOs* and *aao3S*s, grown with or without those stress conditions (Figs. 5D, 6D), led us to examine the reason/s for such phenomenon. It is possible that the absence of AAO2 likely leads to higher availability of MoCo and/or less competition for MoCo sulfuration, resulting in increased activity in the upper activity band that includes AAO3 and AAO1 in WT and aao*2 KOs* and only AAO3 in the *aao3Ss*.

In-vitro sulfuration of AO by employing Na_2_S under anaerobic reducing conditions was previously demonstrated in leaves and roots of WT, recovering significantly the AO activity in the tomato Moco sulfurase (*flacca*) mutant roots and leaves (Sagi et al. 1999), as well as in *aba3* mutant, by employing L-Cysteine (Bittner et al., 2001; Mendel, 2022). Similarly, in-vitro sulfuration was applied and in-gel activity assay of AAO using *trans*-2-nonenal as the substrate was carried out to examine AAO activity levels in the control and the sulfurated extracts of WT, *aao2KO* and *aba3* leaves. Interestingly the uppermost activity band, containing enzyme activities of AAO1 and AAO3 (Nurbekova et al. 2021) was enhanced by 40% in control treated *aao2 KO* mutant as compared to WT’s most upper activity band. Notably, in-vitro sulfuration of WT extracts resulted in a significant increase by 62% in AAO2 activity as well as more than 2-fold enhancement in the heterodimer middle activity band containing a monomer of AAO2, while the uppermost activity band exhibited a slightly lower activity level than the control when *trans-2-*nonenal was used as the substrate. In contrast, the in-vitro sulfuration in *aao2KO* was almost 2-fold higher than in the sulfurated WT and 41% higher than the activity in the *aao2KO* control extract (Supplemental Fig. S6A). These results indicate that AAO2, as the homodimer (lowest migrated activity band) as well as the heterodimer (middle migrated activity band) targets the majority of the sulfuration as compared to the upper activity band (Supplemental Fig. S6A). In support of this notion, employing in-vitro sulfuration in *aba3* mutant that lacks AAO activities led to more than 2-fold higher intensity of AAO2 activity as compared to the heterodimer and the most upper activity bands of the sulfurated extract in *aba3* mutant [calculated using the intensities of the activity bands in *aba3* Supplemental Fig. S6A)]. Importantly, the results indicated that the artificial sulfuration process used partially but significantly recovered AAO activities, as it was ca 60%, 90% and 50% in the upper, middle, and the lowest activity band as compared to the non-sulfurated extract of WT leaves, respectively (Supplemental Fig. S6A).

ABal that was shown as the specific substrate for AAO3 among the four aldehyde oxidases in *Arabidopsis* (Akaba et al. 1998, 1999; Koiwai et al. 2004; Koshiba et al. 1996; Seo et al. 2000a, b) was used to identify the responses of AAO3 to the sulfuration process, as one of the 2 enzymes responsible for the most upper activity band. Notably, while the sulfuration process recovered a faint AAO3 activity band in sulfurated *aba3* that was ca 11% of the activity detected in non-sulfurated WT under control conditions, the sulfuration of *aba3* after UV-C irradiation enhanced AAO3 activity by more than 5-fold (Supplemental Fig. S6B), indicating that UV-C irradiation most likely enhanced the level of inactive unsulfurated AAO3 in *aba3* mutants. Since UV-C irradiation and environmental stress such as Rose Bengal spray were shown to enhance the relative expression of *ABA3* transcription in WT and *aao2 KOs* (Supplemental Fig. S7A and B) as well as the expression of *AAO3* transcript in WT but not in *aao2 KO* mutants (Supplemental Fig. S8) it is likely that UV-C will enhance the levels of active AAO3, at least in the *aao2 KO* mutants mainly by sulfuration of inactive AAO3 proteins.

While control treated plants revealed 2.5-fold higher AAO3 activity in *aao2 KO* mutant compared to WT, the sulfuration process enhanced AAO3 activity in WT by 89% and in *aao2 KO* by 37%, the latter still being higher than WT by 81%. Treating the control WT and *aao2 KO* plants with UV-C irradiation resulted in higher AAO3 activity increase than the sulfuration process, as the activity was enhanced by more than 4-fold in WT and more than 2-fold in *aao2 KO*, being still higher by 11% in the latter (Supplemental Fig. S6B). Notably, trying to enhance AAO3 activity by applying the sulfuration process to UV-C treated plants revealed 18% increased activity in WT and 14% in *aao2 KO*, the latter was still higher by 8% than in WT. These results indicate that while environmental stresses such as UV-C irradiation likely enhance the level of active AAO3 in both WT and the *aao2KO*, the active AAO2 in WT employs a sulfuration process essential for AAO3 activation, resulting in higher availability of activation by sulfuration of the AAO3 in *aao2 KO* compared to AAO3 in WT.

### Summary

The current study demonstrates that AAO2 activity level oppositely affects AAO3 activity, while mutation of AAO2 expression better delays plant senescence by AAO3-enhanced aldehyde detoxification activity in plants exposed to UV-C irradiation or Ros-Bengal spray (Figs. 1C-6). Firstly, it was shown that *aao2KO* can detoxify an array of reactive aldehydes by higher activity than in WT of AAO1 or AAO3 or both exposed to UV-C irradiation or Ros Bengal spray. Additionally, it was shown that AAO3 and/or AAO1 activity increase was followed by AAO2 activity decrease in WT, *aao1KO* and *aao3KO* plants (Figs. 3-4). Secondly, an essential interplay between AAO2 and AAO3 in the regulation of senescence level and carbonyl aldehyde detoxification under the stress conditions was presented. Notably, the delayed senescence and the lower toxic aldehyde levels in *aao2KO* and *aao3Ss* mutants, but not in WT, as well as in *aao1S* and *AAO1OE* mutants treated with UV-C or Rose Bengal, demonstrate the essentiality of AAO2 impairment to plant senescence delay by enhancing aldehyde detoxification activity by AAO3 (Figs.5-6). Finally, employing the sulfuration process (known to activate aldehyde oxidases) in wild-type, *aao2KO*, and *molybdenum-cofactor sulfurase* (*aba3-1*) mutant plants revealed that the active AAO2 in WT employs sulfuration processes essential for AAO3 activity level, resulting in the lower AAO3 activity in WT than AAO3 activity in *aao2KO* (Supplemental Fig.6A and B). These results demonstrate the essential role of AAO2 expression level in regulating AAO3 activity affecting senescence delay in plant leaves by aldehyde detoxification.

## EXPERIMENTAL PROCEDURES

### Chemicals

Acrolein, benzaldehyde, cinnamaldehyde, citral, crotonaldehyde, decanal, dodecanal, heptaldehyde, hexanal, *trans*-2-heptenal, 4-hydroxynonenal-dimethylacetal (HNE-DMA), indole-3-carboxaldehyde, malondialdehyde [tetrabutylammonium salt], 1-naphthaldehyde, nonanal, *trans*-2-nonenal, propionaldehyde, salicylaldehyde, *trans*-2-pentanal, *trans*-2-hexanal, vanillin, diphenyltetrazolium bromide (MTT), phenazine methosulfate (PMS), were purchased from Sigma Aldrich Chemical Company Inc. Abscisic aldehyde was purchased from Toronto Research Chemicals (www.trc-canada.com).

### Plant material and growth conditions

*Arabidopsis thaliana* var. Columbia wild type (WT), T-DNA homozygous knock out (KO) lines for *AAO3* (At2g27150), SAIL_78_H09 (KO9), *AAO1* (At5g20960, SALK_018100), and *AAO2*(At3g43600), KOs SALK_104895 (KO-95) and SAIL_563_G09 (KO-563) were procured from ABRC (http://abrc.osu.edu/). *aba3-1* (At2g16540) used in this study used by us before (Bekturova et al. 2021; Sagi et al. 2002) was procured from the same source. Homozygous lines of T-DNA KOs were screened using PCR with the specific sets of primers (Supplemental Fig. S1A and Supplemental Table *S*1A).

*aao1S* (*aao1* single) mutant was generated by silencing AAO3 in the background of *aao2* (KO-95) and *aao3S* was generated by silencing AAO1 in the background of *aao2* and silencing and AAO2 in the background of *aao1*. The expression of AAOs genes were quantified by qPCR (Supplemental Fig. S2) in *aao1S*, *aao2S* and *aao3S* mutants by employing forward and reverse primers localized to either side of the T-DNA insertion.

For the gene silencing we used the RNA interference system as done before (Brychkova et al. 2007). The sense fragment was ligated into pRNA69 plasmid via the restriction sites XhoI and EcoRI, and the antisense fragment through the restriction sites BamHI and XbaI. The resulting construct was digested, and the fragment containing the 35S promoter and the inserted AAO fragments was inserted via Not1 flanking sites of the binary vector pML-BART.

The generation of AAO1, AAO2 and AAO3 fragments for ligation into pRNA69 plasmid were generated as follows: Forward and reverse primers for the sense fragments were attctcgagCGAGTTCTTCGGATCTTTA and gccgaattcAACATTCTGATCAAAACCA, attCTCGAGTCTTGAGGCTCATGAT and attCTCGAGTCTTGAGGCTCATGAT, attCTCGAGGAAAGAGAAGGTTGATAAT and aggGAATTCTGTTATGAAGCTCAG for for AAO1, AAO2 and AAO3 respectively.

Forward and reverse primers for the antisense fragments were GCCTCTAGACGAGTTCTTCGGATCTTTA and ATAGGATCCAACATTCTGATCAAAACCA, attTCTAGATCTTGAGGCTCATGAT and aggGGATCCTTCAGAGTTTGTTTCA, attTCTAGAGAAAGAGAAGGTTGATAAT and aggGGATCCTGTTATGAAGCTCAG for AAO1, AAO2 and AAO3 respectively. The construct was introduced into Agrobacterium tumefaciens strain GV3101 as described before (Brychkova et al. 2007 and Srivastava et al. 2017) and transformed into the KO mutants using the floral dip method (Clough and Bent,1998). Transformed *Arabidopsis* lines were first selected by resistance to Basta® (glufosinate ammonium: Aventis CropScience https://www.bloomberg.com) and verified by transcript expression level and sequencing.

Seeds were sown in Petri dishes containing half-strength MS medium supplemented with 0.5% plant Agar (https://www.duchefa-biochemie.com/) and were placed at 4°C for 2 days for stratification and then were grown at 14h light/10h darkness in a growth room, 22°C, with 75–85% relative humidity, and 100-150 µmol m^-2^ sec^-1^ as described in (Brychkova et al. 2007). Seven days post germination (DPG), the plants were transferred to 90 mm plates containing half-strength MS medium supplemented with 0.5% sucrose and 1% agar. The biomass of the rosette leaves was determined 3 or 4 days after UV-C irradiation or 17 hours after spraying plants with RB. Untreated plants were used as the control.

### RNA isolation, cDNA preparation, and real-time PCR

To quantify the transcripts using quantitative reverse transcriptase-polymerase chain reaction (qPCR), total RNA was prepared by using the Aurum^TM^ total RNA Mini Kit (Bio-Rad, Hercules, CA) according to the manufacturer’s instructions. The cDNAs were prepared in a 10μl volume containing 350 ng of plant total RNA that was reverse-transcribed with an iScript^TM^ cDNA Synthesis Kit using modified MMLV-derived reverse transcriptase (Bio-Rad), a blend of oligo-d(T) and random hexamer primers according to the manufacturer’s instructions. The generated cDNA was diluted 10 times and the quantitative analysis of transcripts was performed employing the sets of primers enlisted in Supplemental Table S2A as described before (Brychkova et al. 2007; Oshanova et al. 2021).

### DNA sequence analysis

Sequence analysis was performed with the ABI Prism Big Dye Terminator Cycle Sequencing Ready Reaction Kit on an ABI Prism 310 cycle sequencer (PE Applied Biosystems, Warrington, UK).

### Identification of unique peptides in activity bands

To confirm the identity of the AAOs involved in the formation of the lower (most migrated) bands in WT, *aao1* and *aao3* KO, during aldehyde oxidation (Fig. 1C), the activity bands were excised from the native gel, and fractionated with 12% SDS-PAGE. Thereafter the proteins were stained by Coomassie Brilliant Blue, and the individual lanes harboring stained bands were cut from the gel and sent for peptide sequencing at The Smoler Protein Research Center (https://proteomics.net.technion.ac.il/). The proteins were trypsinized and the resulting peptides were separated by HPLC and analyzed by LC-S/MS on Q-Exactive (Thermo Fisher Scientific). Data analysis was done as described in (Kurmanbayeva et al. 2017, Nurbekova et al. 2021).

### Protein extraction and fractionation

Soluble proteins were extracted from *Arabidopsis* rosette leaves as described before (Sagi et al. 1998). Concentrations of total soluble protein in the resulting supernatant were determined according to (Bradford et al. 1976). Native PAGE was carried out as follows; samples were subjected to a Bio-Rad Mini-Protean III slab cell (Bio-Rad, Richmond, CA, USA), with a discontinuous buffer system (Laemmli et.al 1970) in 7.5% (w/v) polyacrylamide separating gels and 4% (w/v) stacking gels in the absence of SDS at 4°C. Tissue/buffer ratio was 1:3 (weight/volume). Extracted crude protein samples were centrifuged at 15000g for 20 min at 4°C and then loaded to the gel after equalization of the proteins. The native-PAGE was carried out using 1.5 mm thick slabs loaded with indicated levels of proteins.

### AAO In-gel activity

AAO activity was tested using different aldehydes in 50mM Tris-HCl, pH 7.5, containing 1 mM 3-(4,5-dimethylthiazol-2-yl)-2,5-diphenyltetrazolium bromide (MTT), 0.1 mM phenazine methosulfate (PMS) and one of various aldehydes (conc. is mentioned at the required places). The reactions were stopped by immersion of the gels in 5% acetic acid. The quantity of resulting formazan activity bands was directly proportional to enzyme activity during a given incubation time, in the presence of excess substrate and tetrazolium salt (Rothe et al. 1974). The gels were scanned, and the intensity of bands was determined using ImageJ software (http://imagej.nih.gov/ij/).

### In vitro sulfuration of crude extracts

In vitro sulfuration was carried out as previously described (Sagi et al. 1999; Sagi et al. 2002) with slight modifications. Protein extraction of rosette leaves was done as described above in the materials and methods section. The high heat stability of plant MoCo-containing hydroxylases in plant leaves (Bower et al. 1978; Omarov et al. 1999; Sagi et al. 1999) allowed to heat the resulted supernatant for 90 sec at 65 °C and additionally centrifuge for 10 min at 4°C as done previously (Nurbekova et al. 2021). Five hundred µl supernatant of the leave extracts as well as one ml of extraction buffer passed through G-25 Sephadex that was previously equilibrated with 6 ml extraction buffer and 1 ml of the desalted protein was collected. For the sulfuration process 450 µl of the desalted protein incubated at room temperature for 40 min together with 20.25 µl 0.1M dithionite, 17 µl of 0.5 M Na_2_S, and 22.5 µl of 1.25 mM methyl viologen was the indicator of the reducing conditions, while gently flushing N_2_ through the mixture to maintain anaerobic conditions. A volume of 59.75 µl extraction buffer was added to 450 µl desalted protein that served as the control. Proteins were equalized before and after sulfuration and AO activity was detected in polyacrylamide gels by stanning after native electrophoresis as previously described. Detection of AO activity was carried out with desalted proteins extracted from WT, *aao2KO* and *aba3-1* mutant leaves.

### UV-C treatment

UV-C treatment was applied by subjecting equal age plants (21 to 23 DPG) to 250 mJ UV-C irradiation during 90 sec, using CL 508 Crosslinkers (UVITec Ltd, UK). WT, *aao2 (*KO-95 and KO-563), *aao1S(aao1S-11)* and *aao3S1(aao3S-1, aao3S-12, aao3S-18)* mutant plants, grown in 1% agar plates containing half-strength MS media and 0.5% sucrose. UV-C irradiation-treated and untreated (control) plants, were placed in the controlled growth room. Photographs of the plants were taken 3 or 4 days after UV-C irradiation and the remaining plants were frozen in liquid N and then stored at −80 °C for further use.

### Rose Bengal treatment in rosette leaves

Twenty-three-day old *aao2* KOs SALK_104895 (KO-95) and SAIL_563_G09 (KO-563), WT, *aao1S (aao1S-11)* and *aao3S2 (aao3S-7, aao3S-12, aao3S-18)* plants grown 3 plants per 90 mm plate were used for Rose Bengal treatment. Rose Bengal was dissolved using a magnetic stirrer in DDW containing 0.01 % Silwet L-77. Fifty µM Rose Bengal was sprayed onto the plates with 3 plants and the remaining solution was removed carefully, and plates were resealed with surgical tape and kept 1 hour under the dark and then were transferred to the normal growth room condition. DDW containing 0.01 % Silwet L-77 was used as the control. Plants were sampled 17 h after Rose Bengal application. The effect of Rose Bengal treatment was documented 72 h after the Rose Bengal application unless mentioned otherwise.

### Chlorophyll and leaf-damage level determination

Total chlorophyll content was measured in extracts of the rosette leaves as described before (Brychkova et al. 2008). The values for remaining chlorophyll content in rosettes were determined as the quantity of chlorophyll per 20 mg for each sample divided by the quantity of chlorophyll per 20 mg mock (control untreated) and were expressed as remaining chlorophyll (%). The severity of leaf damage after Rose Bengal treatment was as follows: 0, no damage; 1, 1-5%; 2, 6-10%; 3, 11–25%; 4, 26-40%; 5, >41% of leaf area damaged. The average leaf damage was then multiplied by the total number of damaged leaves to determine the level of damage.

### Determination and quantification of aldehydes, determination of aldehyde toxicity and ABA determination

Aldehydes were determined according to Matsui et al. 2009 and Mano and Biswas, 2018 as shown recently (Bekturova et al. 2021, Nurbekova et al. 2021). Rosette leaves (250mg) were immersed in 2.5 ml acetonitrile (all acetonitrile mentioned is of HPLC Ultra Gradient Solvent) containing 25 nmol 2-ethylhexanal (as an internal standard) and 0.005% (w/v) butylhydroxytoluene (BHT). The leaves were incubated in a screw-capped glass tube at 60°C for 30 min and the supernatant was collected by decantation. For derivatization of aldehydes, 2,4-dinitrophenylhydrazine (DNPH) was purified (recrystallized) as described before (Mano and Biswas, 2018). Briefly, an equal of commercial DNPH was dissolved in 50 ml of warm (60 °C) acetonitrile. Complete dissolved DNPH was chilled gradually (for several hours) and generated crystals were collected through filter paper. DNPH crystals were dissolved in acetonitrile to make 20 mM solution. To derivatize aldehydes with DNPH, 62.5µl of 20mM DNPH (in acetonitrile) and 48.4µl of 99% formic acid (LC-MS quality) were added to the resulting supernatant and incubated at 25°C for 60 min. Thereafter 2.5ml NaCl (5M) and 450mg NaHCO_3_ were added to the mixture for neutralizing formic acid and mixture was shaken at intervals for 10 min. After centrifugation, the upper layer was collected and dried in vacuum and then dissolved in 250 μl acetonitrile. This solution was applied on a Bond Elute C18 cartridge [sorbent mass 200 mg (Agilent Technologies, USA)], which has been pre-washed with 2 ml acetonitrile, and the pass-through (250 μl) and subsequent wash (150 μl acetonitrile) were combined as the eluted solution. Fifteen µl of the eluted solution were analyzed using Thermo Scientific DionexUltiMate 3000 UHPLC system (http://www.dionex.com/en-us/products/liquid-chromatography/lc-systems/lp-87043.html) with Variable Wavelength VWD-3100 detector. The fifteen µl eluted solution was injected into a Wakosil DNPH-II column (4.6 mm x 150 mm(W), No.17193, Wako Pure Chemical Industries, Ltd., Osaka, Japan, http://www.wako-chem.co.jp/english) with the following elution conditions: 1 ml min-1 flow rate, 0–5 min, 100% HPLC quality Eluent A (Wako Pure Chemical Industries) 5–20 min, a linear gradient from 100% A to 100% HPLC quality Eluent B (Wako Pure Chemical Industries); 20–25 min, 100% B, 25–45 min, a linear gradient from 100% B to 100% C (acetonitrile). The detection wavelength was 340 nm, and the column temperature was 35°C. Aldehydes were identified by their retention times and the retention times of the detected aldehydes were compared with aldehyde standards in the same run. The content of the aldehydes was calculated from the peak area of the corresponding DNP-carbonyl by using appropriate software [CHROMELEON7 (https://www.thermofisher.com)]. Aldehyde toxicity was examined with 21-day old *aao2* KOs and WT plants as done by us before (Nurbekova at al. 2021). The effect of aldehyde application was photographed as mentioned in the figure legends. ABA concentrations in leaves were determined as previously described (Turečková et al. 2009).

### Preparation of DNP-aldehyde standards

DNP-aldehyde standards were prepared as previously described (Mano and Biswass., 2018). 1 mM of DNPH (ca. 0.3 g) was dissolved in 20 ml acetonitrile (HPLC Ultra Gradient Solvent). To the dissolved DNPH solution 1 mM of aldehyde and few drops of formic acid were added. The mixture was kept under a chemical hood by stirring. For acetals, such as acrolein, 1 mmol of aldehyde was dissolved in 30 ml of DNPH solution (5 mM in 2 M HCL). After DNP-Aldehyde precipitation, crystals of DNP-aldehyde standards were collected and dried. The mol amount of DNP-derivative is determined by the weight.

## ACCESSION NUMBERS

Sequence data used in this work can be found in the GenBank/ EMBL data libraries under accession numbers At2g27150 (AAO3), At3g43600 (AAO2), At5g20960 (AAO1).

## ACKNOWLEDGMENTS

M.S. thankfully acknowledge a grant from the Israel Center of Research Excellence (Plant Adaptation grant no. 757/12), and M.St. and V.T. thankfully acknowledge a grant from the European Regional Development Fund-Project (No. CZ.02.1.01/0.0/0.0/16_019/0000827).

## CONFLICT OF INTEREST

All authors have no conflicts of interest to declare.

## AUTHOR CONTRIBUTIONS

Zh.N. participated in designing the research plans and performed the experiments and analysis; S.S. read and commented on the manuscript; ZD.N. participated in in-gel activity analysis, V.T. and M.St. performed the detection of ABA level; (R.O.)read and commented on the manuscript; M.S. conceived the original idea, designed the research plan and supervised the research work. The article was jointly written by Zh. N and M.S.

## SUPPORTING INFORMATION

The following supplemental materials are available in the online version of this article

**Supplemental Figure S1.** Relative transcript expression of AAO2 (At3g43600) and Aldehyde oxidases (AAOs) activity in control and UV-C treated WT, *aao2* [SALK_104895 (KO-95)], *aao1* [At5g20960, SALK_018100, (a1-100)] and *aao3* [At2g27150, SAIL_78_H09 (KO9)] mutant plants.

**Supplemental Figure S2.** Relative transcript expression of AAO1(At5g20960), AAO2 (At3g43600) and AAO3 (At2g27150) in rosette leaves of 23-days post germination *Arabidopsis* WT, *aao1Single (aao1S)* and independent *aao3Singles (aao3S)* mutant plants.

**Supplemental Figure S3.** Determination of UV-C-irradiation-induced senescence and senescence-related factors in rosette leaves of *Arabidopsis* wild-type (WT) and AAO1 overexpression (AAO1OE) mutant plants.

**Supplemental Figure S4.** The effect of exogenously applied benzaldehyde on rosette leaves of wild-type (WT) and *aao2* KO [SALK_104895 (KO-95) and SAIL_563_G09 (KO-563)] mutant plants.

**Supplemental Figure S5.** The effect of UV-C irradiation on the transcript expression of *Nine-cis-epoxy carotenoid dioxygenase 3 (NCED3-2;* At3g14440) gene in rosette leaves of *Arabidopsis* WT and *aao2 KO* [SALK_104895 (KO-95) and SAIL_563_G09 (KO-563)] mutant plants.

**Supplemental Figure S6.** The effect of molybdenum cofactor sulfuration and UV-C irradiation on Aldehyde oxidases (AAOs) activity in WT, *aao2* [SALK_104895 (KO-95)] and the molybdenum cofactor sulfurase (*aba3-1,* At1g16540) mutant plants.

**Supplemental Figure S7.** The effect of UV-C irradiation or Rose Bengal treatment on the transcript expression of ABA3 (At1g16540) in rosette leaves of *Arabidopsis* WT and *aao2* [SALK_104895 (KO-95) and SAIL_563_G09 (KO-563)] mutant plants.

**Supplemental Figure S8.** The effect of UV-C irradiation or Ros Bengal treatment on relative transcript expression of AAO3 (At2g27150) in rosette leaves of *Arabidopsis* WT, *aao2* [SALK_104895 (KO-95) and SAIL_563_G09 (KO-563)] mutant plants.

**Supplemental Table S1.** Verification of AAO2 (At3g43600) KO mutants (KO-95 and KO-563) homozygosity.

**Supplemental Table S2. A.** Details of primers used to carry out quantitative real time PCR analysis and **(B)** results of sequence analysis of the resulting product.

**Supplemental Table S3.** Unique peptides of *Arabidopsis* aldehyde oxidases (AAOs) identified by LC-MS.

## Supplemental Figures

Supplemental **Figure S1.** Relative transcript expression of AAO2 (At3g43600) and Aldehyde oxidases (AAOs) activity in control and UV-C treated WT, *aao2* [SALK_104895 (KO-95)], *aao1* [At5g20960, SALK_018100, (a1-100)] and *aao3* [At2g27150, SAIL_78_H09 (KO9)] mutant plants. A. Transcript expression levels of AAO2 gene in rosette leaves of WT, *aao2 (*KO-95*), aao1(*a1-100) and *aao3* (KO9) 72 hours after UV-C irradiation treatment (red bars) or untreated (blue bars) were compared with the corresponding transcript in WT after normalization to *EF-1a* (At5g60390) transcript as the housekeeping gene and presented as relative expression. Different letters above the bar indicate significant differences (Tukey-Kramer HSD test, P<0.05). Asterisk shows significant differences between treatments within the same genotype (Student’s t test, P < 0.05). B. Aldehyde oxidases (AAOs) in gel activity in 23 days old WT, *aao1 (*a1-100) and *aao3* (KO9) mutant plants. 150 µg crude protein extract from the rosette leaves of each genotype was fractionated by NATIVE PAGE for the in-gel activity assay in a solution containing 100 mM Tris-HCl (pH 7.5), 1 mM 3-(4,5-dimethylthiazol-2-yl)-2,5-diphenyltetrazolium bromide, 0.1 mM phenazine methosulfate and 1 mM acrolein as substrate. The gels were scanned after 2h, and intensity of the activity bands was estimated using ImageJ software (http://imagej.nih.gov/ij/). Each of the obtained intensities was compared with that obtained with UV-C untreated (control) WT (employed as 100%) and presented as relative intensity (RI).

Supplemental **Figure S2.** Relative transcript expression of AAO1(At5g20960), AAO2 (At3g43600) and AAO3 (At2g27150) in rosette leaves of 23-days post germination *Arabidopsis* WT, *aao1Single (aao1S)* and independent *aao3Singles (aao3S)* mutant plants. The expression level of each of the transcripts in WT, *aao1S (aao1S-11) and aao3Ss (aao3S-1, aao3S-7, aao3S-11, aao3S-12, aao3S-18)* mutants was compared with the corresponding transcript in WT after normalization to the transcript of *EF-1a (*At5g60390), as the housekeeping gene and presented as relative expression. Different letters above the bar show significant differences (Tukey-Kramer HSD test, P<0.05). *aao1S* was generated by silencing AAO3 in *aao2* [SALK_104895 (KO-95)], and *aao3S* was generated by silencing AAO1 in *aao2(KO-95)* plants or silencing AAO2 in *aao1* [SALK_018100 (a1-100)] plants.

Supplemental **Figure S3.** Determination of UV-C-irradiation-induced senescence and senescence-related factors in rosette leaves of *Arabidopsis* wild-type (WT) and AAO1 overexpression (AAO1OE) mutant plants. **A.** Representative photograph of WT and AAO1OE rosette leaves in response to UV-C irradiation. 23-day post germination (DPG) plants exposed to 250 mJ UV-C irradiation were kept in the growth room for 72 hours and thereafter documented. Scale bar=1 cm. Rosette leaves were collected 3 days after UV-C treatment and rosette leaves of plants that were not exposed to UV-C were used as the control. The first six leaves, oldest to youngest, from left to right are presented. **B.** Remaining chlorophyll in first six leaves (oldest to youngest from left to right) after UV-C treatment. **C.** Indicated aldehyde profiling in control (blue bars) and UV-C treated (red bars) plants. Leaves from 3 different plants were taken as one replica and the bars show the average of at least 4 replicas. Different letters above the bar indicate significant difference (Tukey-Kramer HSD test, P<0.05). Asterisk shows significant differences between treatments within the same genotype (Student’s t test, P < 0.05).

Supplemental **Figure S4.** The effect of exogenously applied benzaldehyde on rosette leaves of wild-type (WT) and *aao2* KO [SALK_104895 (KO-95) and SAIL_563_G09 (KO-563)] mutant plants. **A.** The appearance of WT and *aao2KO* rosette leaves 3 days after exogenously applied 4.5 mM benzaldehyde and water applied as the control. The benzaldehyde solution or water (control) were added to the plants grown on plates (23-day post germination (DPG) plants) for 3 h and thereafter were removed. Arrows indicate damaged area in leaves. Scale bar=1cm. **B.** Remaining chlorophyll detected in the six oldest leaves of benzaldehyde and control treated plants. Different letters above the bars indicate significant differences according to the Tukey-Kramer HSD mean-separation test (P <0.05).

Supplemental **Figure S5.** The effect of UV-C irradiation on the transcript expression of *Nine-cis-epoxy carotenoid dioxygenase 3 (NCED3-2;* At3g14440) gene in rosette leaves of *Arabidopsis* WT and *aao2 KO* [SALK_104895 (KO-95) and SAIL_563_G09 (KO-563)] mutant plants. Controls and UV-C treated are marked with blue and red colors respectively. The *NCED3-2* transcript expression in UV-C irradiation treated or control treated WT and *aao2 KO* (KO-95, KO-563) leaves were compared with the respective transcript in WT control after normalization to the *Arabidopsis EF-1a (*At5g60390) and presented as the relative expression. Values with different letters above the bar are significantly different according to the Turkey-Kramer HSD mean-separation test (P < 0.05). Asterisk shows significant difference between treatments within the same genotype (Student’s t test, P < 0.05).

Supplemental **Figure S6.** The effect of molybdenum cofactor sulfuration and UV-C irradiation on Aldehyde oxidases (AAOs) activity in WT, *aao2* [SALK_104895 (KO-95)] and the molybdenum cofactor sulfurase (*aba3-1,* At1g16540) mutant plants. **A.** AAOs in gel activity of not Sulfurated (NS) and Sulfurated (S) control WT, aao*2* (KO-95) and *aba3* mutant plants using trans-2-nonenal as the substrate. **B.** Aldehyde oxidase 3 in gel activity of NS and S control as well as UV-C treated WT, aao*2* (KO-95) and *aba3* mutant plants with abscisic aldehyde as the specific substrate for AAO3 activity. 150 µg crude protein extracted from UV-C treated or untreated WT, aao*2* (KO-95) and *aba3* rosette leaves that were sulfurated (S) or not sulfurated (NS), was fractionated by NATIVE PAGE for the in-gel activity, assayed for 1h in solution containing 100 mM Tris-HCl (pH 7.5), 1 mM 3-(4,5-dimethylthiazol-2-yl)-2,5-diphenyltetrazolium bromide, 0.1 mM phenazine methosulfate and 1 mM *trans*-2-nonenal or 0.1 mM abscisic aldehyde as the substrate. The gels were scanned, and the intensity of the activity bands was estimated using ImageJ software (http://imagej.nih.gov/ij/) and compared with that obtained with non Sulfurated (NS) WT control (employed as 100%) and presented as relative intensity (RI).

Supplemental **Figure S7.** The effect of UV-C irradiation or Ros Bengal treatment on the transcript expression of ABA3 (At1g16540) in rosette leaves of *Arabidopsis* WT and *aao2* [SALK_104895 (KO-95) and SAIL_563_G09 (KO-563)] mutant plants. **A.** The transcript expression of ABA3 gene in rosette leaves of WT and aao*2* (KO-95, KO-563) 72 hours after UV-C irradiation (red bars) or controls (untreated) (blue bars) were compared with the corresponding transcript in WT after normalization to the transcript of *EF-1a (*At5g60390) as the housekeeping gene and presented as relative expression. **B.** The transcript expression of ABA3 (At1g16540) gene in rosette leaves of WT and *aao2* (KO-95, KO-563) 17 hours after Rose Bengal (RB) treated (red bars) or untreated (blue bars) were compared with the corresponding transcript in WT after normalization to the transcript of *EF-1a (*At5g60390) as the housekeeping gene and presented as relative expression. Different letters above the bar show significant differences (Tukey-Kramer HSD test, P<0.05). Asterisk shows significant differences between treatments within the same genotype (Student’s t test, P < 0.05).

Supplemental **Figure S8.** The effect of UV-C irradiation or Rose Bengal treatment on relative transcript expression of AAO3 (At2g27150) in rosette leaves of *Arabidopsis* WT, *aao2* [SALK_104895 (KO-95) and SAIL_563_G09 (KO-563)] mutant plants. **A.** The transcript expression of AAO3 in rosette leaves of *Arabidopsis* WT, *aao2* (KO-95, KO-563) 72 hours after UV-C irradiation (red bars) or in controls [untreated (blue bars)] were compared with the corresponding transcript in WT after normalization to the transcript of *EF-1a* (At5g60390), as the housekeeping gene and presented as relative expression. **B.** The transcript expression levels of AAO3 gene in rosette leaves KO-95, KO-563 17 hours after Rose Bengal (RB) treated (red bars) or untreated (blue bars) were compared with the corresponding transcript in WT after normalization to the transcript of *EF-1a (*At5g60390) as the housekeeping gene and presented as relative expression. Different letters above the bar show significant differences (Tukey-Kramer HSD test, P<0.05). Asterisk shows significant differences between treatments within the same genotype (Student’s t test, P < 0.05).

**Supplemental Tables**

Supplemental **Table S1.** Verification of AAO2 (At3g43600) KO mutants (KO-95 and KO-563) homozygosity. (A) List of primers used for verification of AAO2 KO (SALK_-_104895.22.00.n and SAIL_563_G09) and mutants. (B) *aao2* KO mutants’ identity, chromosomes position and amplicons length.

Supplemental **Table S2.** (A) Details of primers used to carry out quantitative real time PCR analysis and (B) the results of sequence analysis of the resulting product of AAO2 and aba3. The Sequence of *Arabidopsis Aldehyde oxidase 3 (AAO3; At2g27150), Arabidopsis Aldehyde oxidase 1 (AAO1; At5g20960), Nine-cis-epoxy carotenoid dioxygenase 3 (NCED3-2; At3g14440)* was already shown in Nurbekova et.al.,2021 and *Elongation factor 1-alpha (EF1α; AT5G60390)* was shown in Srivastava et.al., 2017.

Supplemental **Table S3.** Unique peptides of *Arabidopsis* aldehyde oxidases (AAOs) identified by LC-MS. In gel activity was conducted as shown in Figure 3D and lower bands from the indicated genotypes were excised followed by their peptide sequencing.

## Notes

### Competing Interest Statement

The authors have declared no competing interest.

### Summary of Updates

To add supplemental Figures and Tabled not uploaded before.

