## Supplementary material for "AAO2 impairment improves aldehyde detoxification by AAO3 in Arabidopsis leaves exposed to UVC or Rose Bengal": Supplanental Figures and Tables

**A**

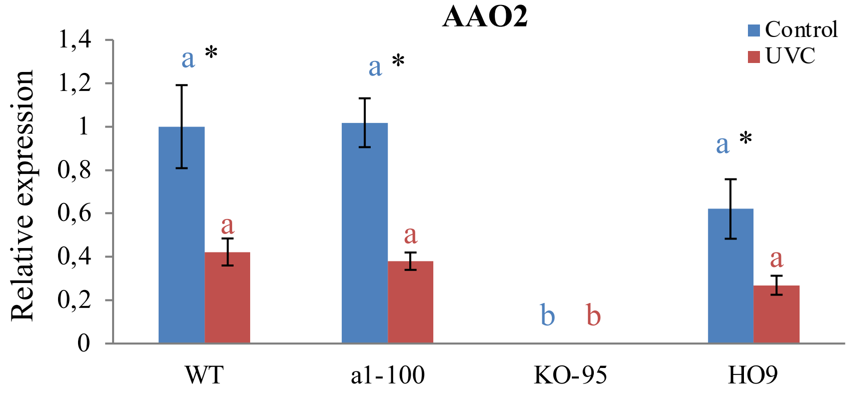


**B**

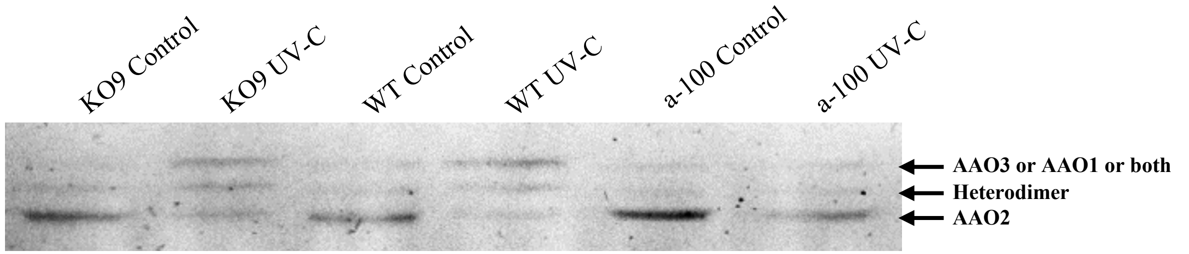


| | | | | | | | |
|---|---|---|---|---|---|---|---|
| RI | 74 | 238 | 100 | 327 | 84 | 91 | ← **AAO3 or AAO1 or both** |
| RI | 117 | 158 | 100 | 143 | 57 | 74 | ← **Heterodimer** |
| RI | 115 | 41 | 100 | 26 | 140 | 63 | ← **AAO2** |

acrolein as the substrate

**Supplemental Figure S1.** Relative transcript expression of AAO2 (At3g43600) and Aldehyde oxidases (AAOs) activity in control and UV-C treated WT, *aao2* [SALK_104895 (KO-95)], *aao1* [At5g20960, SALK_018100, (a1-100)] and *aao3* [At2g27150, SAIL_78_H09 (KO9)] mutant plants**. A.** Transcript expression levels of AAO2 gene in rosette leaves of WT, *aao2 (*KO-95*), aao1(*a1-100*)* and *aao3* (KO9) 72 hours after UV-C irradiation treatment (red bars) or untreated (blue bars) were compared with the corresponding transcript in WT after normalization to *EF-1a* (At5g60390) transcript as the housekeeping gene and presented as relative expression. Different letters above the bar indicate significant differences (Tukey-Kramer HSD test, P<0.05). Asterisk shows significant differences between treatments within the same genotype (Student's t test, P < 0.05). **B.** Aldehyde oxidases (AAOs) in gel activity in 23 days old WT, *aao1 (*a1-100) and *aao3* (KO9) mutant plants. 150 µg crude protein extract from the rosette leaves of each genotype was fractionated by NATIVE PAGE for the in-gel activity assay in a solution containing 100 mM Tris-HCl (pH 7.5), 1 mM 3-(4,5-dimethylthiazol-2-yl)-2,5-diphenyltetrazolium bromide, 0.1 mM phenazine methosulfate and 1 mM acrolein as substrate. The gels were scanned after 2h, and intensity of the activity bands was estimated using ImageJ software (http://imagej.nih.gov/ij/). Each of the obtained intensities was compared with that obtained with UV-C untreated (control) WT (employed as 100%) and presented as relative intensity (RI).

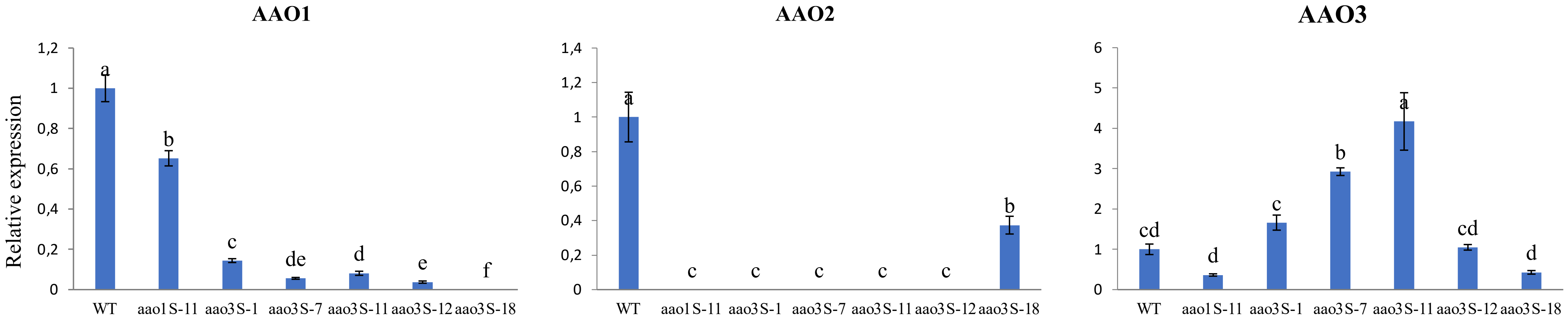


**Supplemental Figure S2.** Relative transcript expression of AAO1(At5g20960), AAO2 (At3g43600) and AAO3 (At2g27150) in rosette leaves of 23- days post germination *Arabidopsis* WT, *aao1Single (aao1S)* and independent *aao3Singles (aao3S)* mutant plants. The expression level of each of the transcripts in WT, *aao1S (aao1S-11) and aao3Ss (aao3S-1, aao3S-7, aao3S-11, aao3S-12, aao3S-18)* mutants was compared with the corresponding transcript in WT after normalization to the transcript of *EF-1a (*At5g60390*)*, as the housekeeping gene and presented as relative expression. Different letters above the bar show significant differences (Tukey-Kramer HSD test, $P<0.05$). *aao1S* was generated by silencing AAO3 in *aao2* [SALK_104895 (KO-95)], and *aao3S* was generated by silencing AAO1 in *aao2(KO-95)* plants or silencing AAO2 in *aao1* [SALK_018100 (a1-100)] plants.

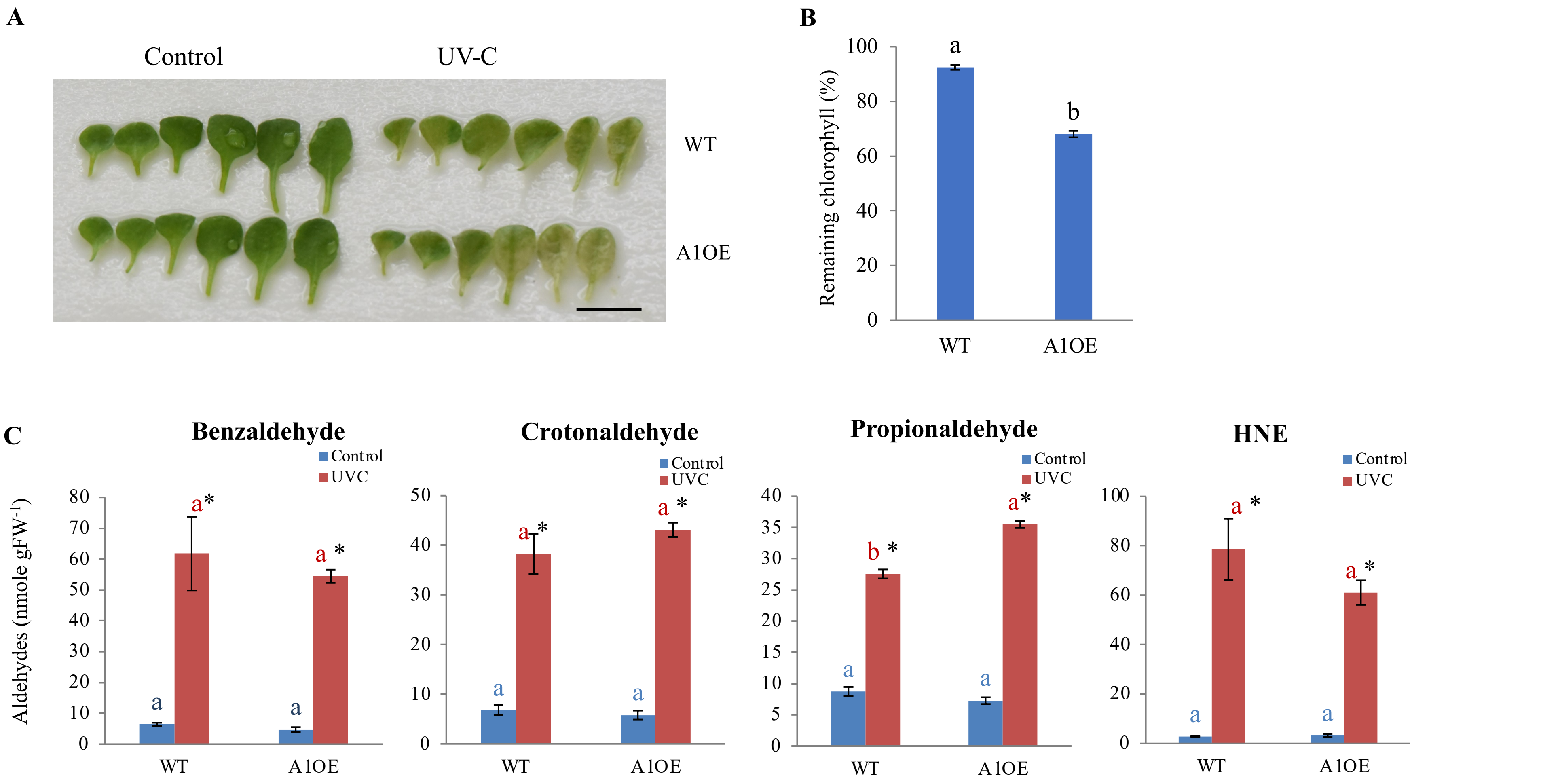


**Supplemental Figure S3.** Determination of UV-C-irradiation-induced senescence and senescence-related factors in rosette leaves of *Arabidopsis* wild-type (WT) and AAO1 overexpression (AAO1OE) mutant plants. **A.** Representative photograph of WT and AAO1OE rosette leaves in response to UV-C irradiation. 23-day post germination (DPG) plants exposed to 250 mJ UV-C irradiation were kept in the growth room for 72 hours and thereafter documented. Scale bar=1 cm. Rosette leaves were collected 3 days after UV-C treatment and rosette leaves of plants that were not exposed to UV-C were used as the control. The first six leaves, oldest to youngest, from left to right are presented. **B.** Remaining chlorophyll in first six leaves (oldest to youngest from left to right) after UV-C treatment. **C.** Indicated aldehyde profiling in control (blue bars) and UV-C treated (red bars) plants. Leaves from 3 different plants were taken as one replica and the bars show the average of at least 4 replicas. Different letters above the bar indicate significant difference (Tukey-Kramer HSD test, $P<0.05$). Asterisk shows significant differences between treatments within the same genotype (Student's t test, $P < 0.05$).

A

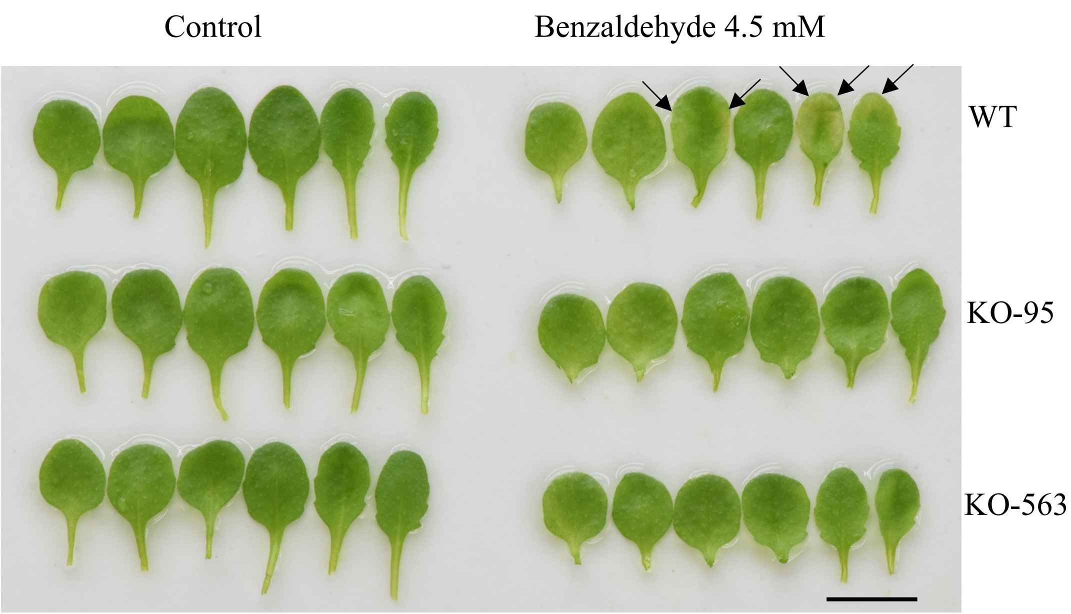


B

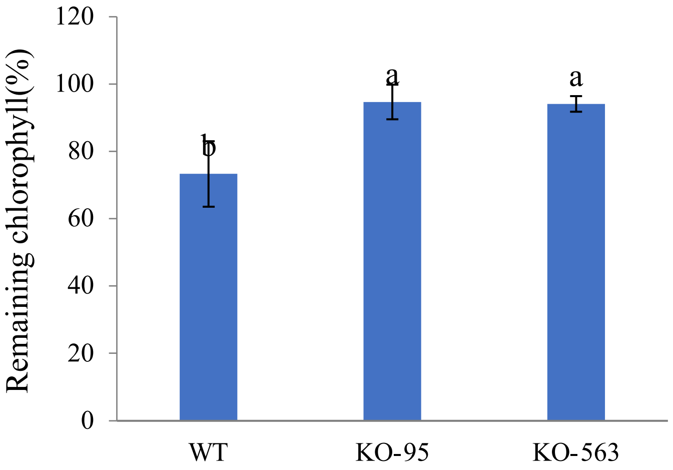


**Supplemental Figure S4**. The effect of exogenously applied benzaldehyde on rosette leaves of wild-type (WT) and *aao2* KO [SALK_104895 (KO-95) and SAIL_563_G09 (KO-563)] mutant plants. **A.** The appearance of WT and *aao2KO* rosette leaves 3 days after exogenously applied 4.5 mM benzaldehyde and water applied as the control. The benzaldehyde solution or water (control) were added to the plants grown on plates (23-day post germination (DPG) plants) for 3 h and thereafter were removed. Arrows indicate damaged area in leaves. Scale bar=1cm. **B.** Remaining chlorophyll detected in the six oldest leaves of benzaldehyde and control treated plants. Different letters above the bars indicate significant differences according to the Tukey-Kramer HSD mean-separation test ($P < 0.05$).

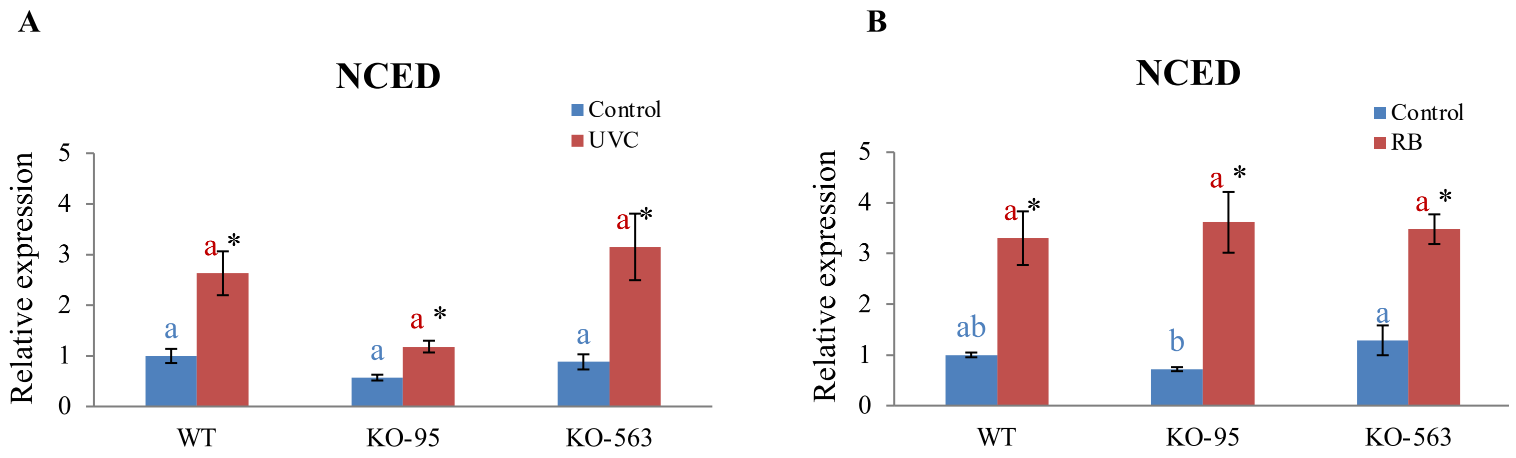


**Supplemental Figure S5.** The effect of UV-C irradiation on the transcript expression of *Nine-cis-epoxy carotenoid dioxygenase 3 (NCED3-2;* At3g14440*)* gene in rosette leaves of *Arabidopsis* WT and *aao2 KO* [SALK_104895 (KO-95) and SAIL_563_G09 (KO-563)] mutant plants. Controls and UV-C treated are marked with blue and red colors respectively. The *NCED3-2* transcript expression in UV-C irradiation treated or control treated WT and *aao2 KO* (KO-95, KO-563) leaves were compared with the respective transcript in WT control after normalization to the *Arabidopsis EF-1a (*At5g60390*)* and presented as the relative expression. Values with different letters above the bar are significantly different according to the Turkey-Kramer HSD mean-separation test ($P < 0.05$). Asterisk shows significant difference between treatments within the same genotype (Student's t test, $P < 0.05$).

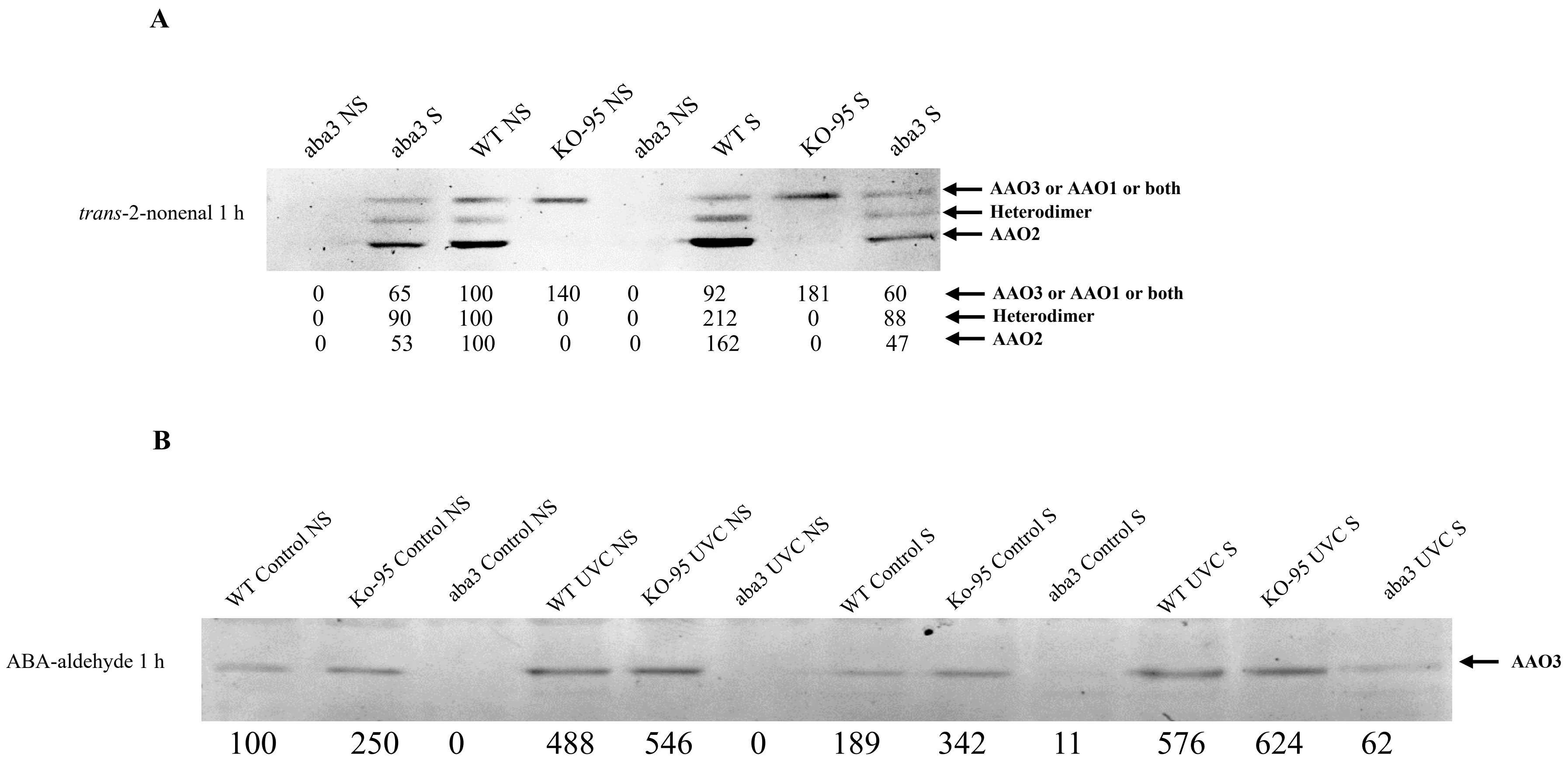


**Supplemental Figure S6.** The effect of molybdenum cofactor sulfuration and UV-C irradiation on Aldehyde oxidases (AAOs) activity in WT, *aao2* [SALK_104895 (KO-95)] and the molybdenum cofactor sulfurase (*aba3-1,* At1g16540) mutant plants. **A.** AAOs in gel activity of not Sulfurated (NS) and Sulfurated (S) control WT, aao*2* (KO-95) and *aba3* mutant plants using trans-2-nonenal as the substrate. **B.** Aldehyde oxidase 3 in gel activity of NS and S control as well as UV-C treated WT, aao*2* (KO-95) and *aba3* mutant plants with abscisic aldehyde as the specific substrate for AAO3 activity. 150 µg crude protein extracted from UV-C treated or untreated WT, aao*2* (KO-95) and *aba3* rosette leaves that were sulfurated (S) or not sulfurated (NS), was fractionated by NATIVE PAGE for the in-gel activity, assayed for 1h in solution containing 100 mM Tris-HCl (pH 7.5), 1 mM 3-(4,5-dimethylthiazol-2-yl)-2,5-diphenyltetrazolium bromide, 0.1 mM phenazine methosulfate and 1 mM *trans*-2-nonenal or 0.1 mM abscisic aldehyde as the substrate. The gels were scanned, and the intensity of the activity bands was estimated using ImageJ software (http://imagej.nih.gov/ij/) and compared with that obtained with non Sulfurated (NS) WT control (employed as 100%) and presented as relative intensity (RI).

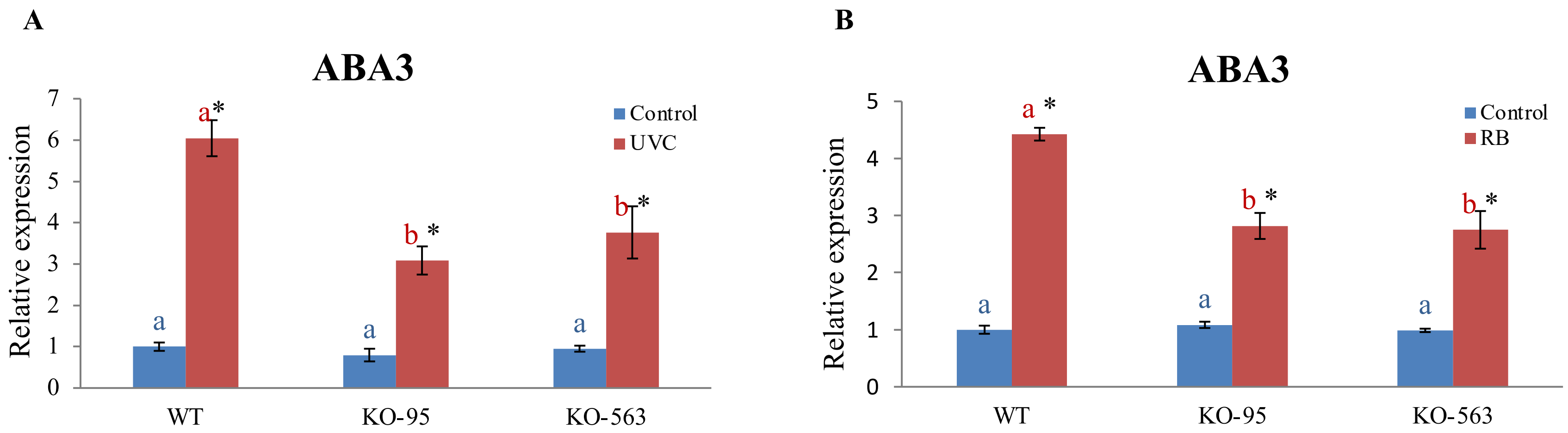


**Supplemental Figure S7.** The effect of UV-C irradiation or Rose Bengal treatment on the transcript expression of ABA3 (At1g16540) in rosette leaves of *Arabidopsis* WT and *aao2* [SALK_104895 (KO-95) and SAIL_563_G09 (KO-563)] mutant plants. **A.** The transcript expression of ABA3 gene in rosette leaves of WT and aao*2* (KO-95, KO-563) 72 hours after UV-C irradiation (red bars) or controls (untreated) (blue bars) were compared with the corresponding transcript in WT after normalization to the transcript of *EF-1a (*At5g60390*)* as the housekeeping gene and presented as relative expression. **B.** The transcript expression of ABA3 (At1g16540) gene in rosette leaves of WT and *aao2* (KO-95, KO-563) 17 hours after Rose Bengal (RB) treated (red bars) or untreated (blue bars) were compared with the corresponding transcript in WT after normalization to the transcript of *EF-1a (*At5g60390*)* as the housekeeping gene and presented as relative expression. Different letters above the bar show significant differences (Tukey-Kramer HSD test, $P<0.05$). Asterisk shows significant differences between treatments within the same genotype (Student's t test, $P < 0.05$).

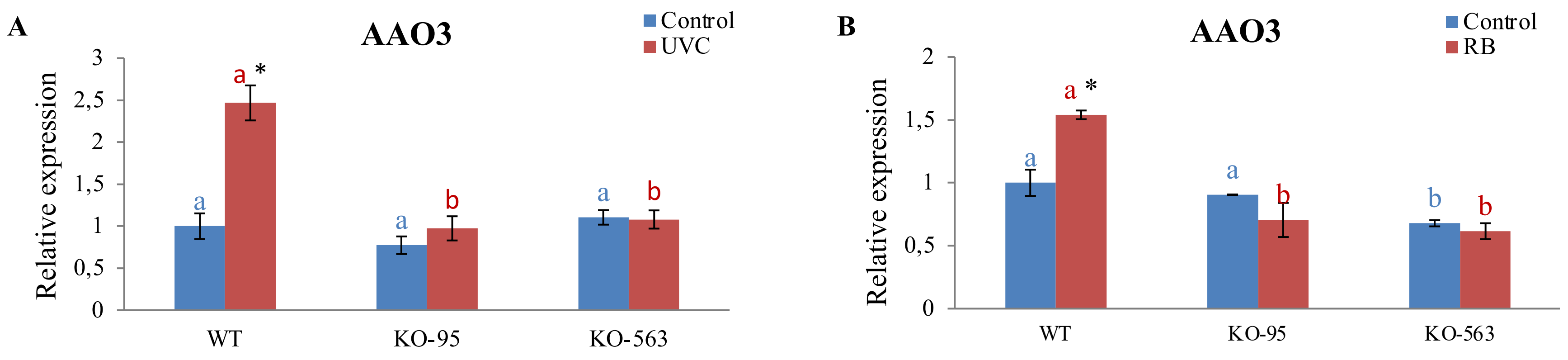


**Supplemental Figure S8.** The effect of UV-C irradiation or Rose Bengal treatment on relative transcript expression of AAO3 (At2g27150) in rosette leaves of *Arabidopsis* WT, *aao2* [SALK_104895 (KO-95) and SAIL_563_G09 (KO-563)] mutant plants. **A.** The transcript expression of AAO3 in rosette leaves of *Arabidopsis* WT, *aao2* (KO-95, KO-563) 72 hours after UV-C irradiation (red bars) or in controls [untreated (blue bars)] were compared with the corresponding transcript in WT after normalization to the transcript of *EF-1a* (At5g60390), as the housekeeping gene and presented as relative expression. **B.** The transcript expression levels of AAO3 gene in rosette leaves KO-95, KO-563 17 hours after Rose Bengal (RB) treated (red bars) or untreated (blue bars) were compared with the corresponding transcript in WT after normalization to the transcript of *EF-1a (*At5g60390*)* as the housekeeping gene and presented as relative expression. Different letters above the bar show significant differences (Tukey-Kramer HSD test, $P<0.05$). Asterisk shows significant differences between treatments within the same genotype (Student's t test, $P < 0.05$).

**Supplemental Table S1.** Verification of AAO2 (At3g43600) KO mutants (KO-95 and KO-563) homozygosity. (A) List of primers used for verification of AAO2 KO (SALK-__104895.22.00.n and SAIL_563_G09 ) and mutants. (B) *aao2* KO mutants' identity, chromosomes position and amplicons length.

**A**

| Name | Oligonucleotide sequence (**5'->3'**) |
|---|---|
| LB1.3 | ATTTTGCCGATTTCGGAAC |
| IB2 | GCTTCCTATTATATCTTCCCAAATTACCAATACA |
| LP (SALK__104895.22.00.n) or (KO-95) | ACTGCATGGGAGTGTCTTTTG |
| RP (SALK__104895.22.00.n) or (KO-95) | GAGGTTTTGGAGGGAAATCTG |
| LP(SAIL_563_G09) or (KO-563) | TGGTCCAAGGAAATTTCACAG |
| RP (SAIL_563_G09) or (KO-563) | GCACAGCGATTAATCTCCAAG |

**B**

| **aao2 mutant identity** | **Insertion location** | **Amplicon length in WT with LP+RP(bp)** | **Amplicon length in mutant with LB+RP(bp)** |
|---|---|---|---|
| SALK_104895.22.00.n or KO-95 | chr3 15514301 | 1236 | 597-897 |
| SAIL_563_G09 or KO-563 | chr3 15513128 | 1162 | 537-837 |

**Supplemental Table S2.** (A) Details of primers used to carry out quantitative real time PCR analysis and (B) the results of sequence analysis of the resulting product of AAO2 and aba3. The Sequence of *Arabidopsis Aldehyde oxidase 3 (AAO3; At2g27150), Arabidopsis Aldehyde oxidase 1 (AAO1; At5g20960), Nine-cis-epoxy carotenoid dioxygenase 3 (NCED3-2; At3g14440)* was already shown in Nurbekova et.al.,2021 and *Elongation factor 1-alpha (EF1α; AT5G60390)* was shown in Srivastava et.al., 2017.

**A.** Details of primers used to carry out quantitative real time PCR analysis.

| **Gen (Identity)** | **Oligonucleotide sequence (5'-)3')** | **Size of amplicon** |
|---|---|---|
| Arabidopsis Aldehyde oxidase 3 (AAO3; At2g27150) | For: GTTGGAGCTGCCTTACAAGCTTCT<br>Rev: CAGTAGGAGTTACATTCTCTCTGAAGC | 151 |
| Arabidopsis Aldehyde oxidase 1 (AAO1; At5g20960) | For: CCACCTATACTGTCCTTAGAAGAAGC<br>Rev: GTGTCTCCATATAGAAGAAGTACTGTG | 175 |
| Arabidopsis Aldehyde oxidase 2 (AAO2; At3g43600 ) | For: GCGTTTTTCATGACATCCACCA<br>Rev: TCCCGCTGTCGATTTAGGACA | 196 |

| | | |
|---|---|---|
| *Arabidopsis thaliana aba deficient3, Low Osmotic Stress 5* (ABA3; At1g16540 ) | For: CTTATTTTAGTGGAGGCACTGTTGCTG<br>Rev: TGTGCATCCAAATTGCAGAAGGTGTAAG | 177 |
| *Elongation factor 1-alpha* (*EF1α*; At5g60390) | For: CAGGACATCGTGATTTCATCAAG<br>Rev: TCCATCTTGTTACAACAGCAAAT | 190 |
| *Nine-cis-epoxy carotenoid dioxygenase 3 (NCED3-2; At3g14440)* | For: TCGAACCGACCAACGGTTTT<br>Rev: GAGCTGCAGCCGGTATAGTCG | 160 |

### *B. 1. Arabidopsis Aldehyde oxidase 2 (AAO2; At3g43600 )*

```
ForwardSeq      ----------------------------------TCCCGCTGTCGATTTAGGACAGATTG
AAO2            CAGATATCTTATATGACTGTGGAAAAAGTCTCAATCCCGCTGTCGATTTAGGACAGATTG
                                                  **************************

ForwardSeq      AAGGATCTTTTGTTCAAGGACTTGGGTTTTTCATGCTTGAAGAGTACATAGAAGATCCAG
AAO2            AAGGATCTTTTGTTCAAGGACTTGGGTTTTTCATGCTTGAAGAGTACATAGAAGATCCAG
                ************************************************************

ForwardSeq      AAGGACTCCTTCTGACGGATAGCACATGGACATACAAGATTCCAACAGT-GACACCATTC
AAO2            AAGGACTCCTTCTGACGGATAGCACATGGACATACAAGATTCCAACAGTTGACACCATTC
                ************************************************* **********


ForwardSeq      ---AACAGTCAAAT--CGAGAA--------------------------------------
AAO2            CTAAACAGTTCAATGTCGAGATACTAAATGGTGGATGTCATGAAAAACGCGTACTCTCTT
                   ******  ***  *****


ReverseSeq      WAACMSSYMTCTGWAAAGGWTWGGGKTTTK--MTGCTTGWAGAGTACATAGAAGATCCAG
AAO2            AAGGATCTTTTGTTCAAGGACTTGGGTTTTTCATGCTTGAAGAGTACATAGAAGATCCAG
                 *       *     ****    ** ***    ****** ********************

ReverseSeq      AAGGACTCCTTCTGACGGATAGCACATGGACATACAAGATTCCAACAGTTGACACCATTC
AAO2            AAGGACTCCTTCTGACGGATAGCACATGGACATACAAGATTCCAACAGTTGACACCATTC
                ************************************************************

ReverseSeq      CTAAACAGTTCAATGTCGAGATACTAAATGGTGGATGTCATGAAAAACGCA---------
AAO2            CTAAACAGTTCAATGTCGAGATACTAAATGGTGGATGTCATGAAAAACGCGTACTCTCTT
                **************************************************
```

### ***2. Arabidopsis thaliana aba3* [*Molybdenum Cofactor Sulfurase* (ABA3; At1g16540)] mutant.**

```
ReverseSeq      ------------------------------------------------------------
aba3            TCTACAAGTTATTTGGTTATCCTACTGGGCTTGGCGCTCTCCTTGTACGGAATGATGCAG


ReverseSeq      -----------------TCTTATTTTAGTGGAGGCACTGTTGCTGCTTCAATTGCTGACA
aba3            CCAAATTGCTCAAAAAGACTTATTTTAGTGGAGGCACTGTTGCTGCTTCAATTGCTGACA
                                  ******************************************

ReverseSeq      TCGACTTTGTAAAAAGAAGGGAAAGGGTGGAGGAGTTTTTTGAGGATGCGTTCTGCTTCA
aba3            TCGACTTTGTAAAAAGAAGGGAAAGGGTGGAGGAGTTTTTTGAGGATG-GTTCTGCTTCA
                ************************************************ ***********

ReverseSeq      TTCCTGAGCATAGCAGCCATCCGTCA-GGCTTCAAATTACTCAAATCTG-----------
```

```
aba3              TTCCTGAGCATAGCAGCCATCCGTCATGGCTTCAAATTACTCAAGTCGCTTACACCTTCT
                  ************************** ***************** **



ForwardSeq        ----------------------------------------------TTCAATTGCTGA-A
aba3              CCAAATTGCTCAAAAAGACTTATTTTAGTGGAGGCACTGTTGCTGCTTCAATTGCTGACA
                                                                ************ *

ForwardSeq        TCG-CTTTGTAAAAAGAAGGGAAAGGGTGGAGGAGTTTTTTGAGGATGGTTCTGCTTCAT
aba3              TCGACTTTGTAAAAAGAAGGGAAAGGGTGGAGGAGTTTTTTGAGGATGGTTCTGCTTCAT
                  *** ********************************************************

ForwardSeq        TCCTGACCATAGCAGCCATCCGTCATGGCTTCAAATTACTCAAGTCGCTTACACCTTCTG
aba3              TCCTGAGCATAGCAGCCATCCGTCATGGCTTCAAATTACTCAAGTCGCTTACACCTTCTG
                  ****** *****************************************************

ForwardSeq        CAATTTGGATGCACAA--------------------------------------------
aba3              CAATTTGGATGCACACAACGTCACTTTCCATATATGTGAAAAAGAAGCTTCAGGCTTTAC
                  ***************
```

**Supplemental Table S3.** Unique peptides of *Arabidopsis* aldehyde oxidases (AAOs) identified by LC-MS. In gel activity was conducted as shown in Figure 3D and lower bands from the indicated genotypes were excised followed by their peptide sequencing.

| WT | *aao1 KO* | *aao3 KO* |
|---|---|---|
| | **AAO2 unique peptides** | |
| VSPGVEK | VGGGFGGK*(1,2,3) | VSPGVEK |
| AVSGNLCR*(1,2) | SVDSGMYR | SSLAPGFLFK |
| SVDSGMYR | IPHLKEIR | DGFHPIHKR |
| IPTVDTIPK*(2,3) | VIAALKEIR | KGDKDSSSLTR |
| SSLAPGFLFK | SSLAPGFLFK | GMAEADHQILSSEIR |
| KGDKDSSSLTR | GMAEADHQILSSEIR | ENQNGVEIGSVVTISK |
| LATHMEMIAAR | ENQNGVEIGSVVTISK | ASGEPPLLLAASVHCATR *(2,3) |
| NFGSIGGNLVMAQR | | LKPLMER |
| GMAEADHQILSSEIR | | VIAALK |
| ENQNGVEIGSVVTISK | | |

* Peptide sequences shares identity with other AAOs [2-AAO2 (Protein ID-BAA28625), 3-AAO3 (AAD22498), 4-AAO4 BAA90299, AAO1 (BAA28624)].
